# A new high-throughput tool to screen mosquito-borne viruses in Zika virus endemic/epidemic areas

**DOI:** 10.1101/764704

**Authors:** Sara Moutailler, Lena Yousfi, Laurence Mousson, Elodie Devillers, Marie Vazeille, Anubis Vega-Rúa, Yvon Perrin, Frédéric Jourdain, Fabrice Chandre, Arnaud Cannet, Sandrine Chantilly, Johana Restrepo, Amandine Guidez, Isabelle Dusfour, Filipe Vieira Santos de Abreu, Taissa Pereira dos Santos, Davy Jiolle, Tessa M. Visser, Constantianus J.M. Koenraadt, Merril Wongsokarijo, Mawlouth Diallo, Diawo Diallo, Alioune Gaye, Sébastien Boyer, Veasna Duong, Géraldine Piorkowski, Christophe Paupy, Ricardo Lourenco de Oliveira, Xavier de Lamballerie, Anna-Bella Failloux

## Abstract

Mosquitoes are vectors of arboviruses affecting animal and human health. Arboviruses circulate primarily within an enzootic cycle and recurrent spillovers contribute to the emergence of human-adapted viruses able to initiate an urban cycle involving anthropophilic mosquitoes. The increasing volume of travel and trade offers multiple opportunities for arbovirus introduction in new regions. This scenario has been exemplified recently with the Zika pandemic. To incriminate a mosquito as vector of a pathogen, several criteria are required such as the detection of natural infections in mosquitoes. In this study, we used a high-throughput chip based on the BioMark^TM^ Dynamic arrays system capable of detecting 64 arboviruses in a single experiment. A total of 17,958 mosquitoes collected in Zika-endemic/epidemic countries (Brazil, French Guiana, Guadeloupe, Suriname, Senegal, and Cambodia) were analyzed. Here we show that this new tool can detect endemic and epidemic viruses in different mosquito species in an epidemic context. Thus, this fast and low-cost method can be suggested as a novel epidemiological surveillance tool to identify circulating arboviruses.

## 1. Introduction

The World Health Organization stated in February 2016 that Zika infection was considered as a public health emergency of international concern (1) opening a new chapter in the history of vector-borne diseases. Arboviruses are viruses transmitted among vertebrate hosts by arthropod vectors. Successful transmission of an arbovirus relies on a complex life cycle in the vector, which after midgut infection and dissemination, is released in saliva for active transmission to the vertebrate host (2). Arboviruses belong to nine families: Asfarviridae, Flaviviridae, Orthomyxoviridae, Reoviridae, Rhabdoviridae, the newly recognized Nyamiviridae (order Mononegavirales) and the families Nairoviridae, Phenuiviridae and Peribunyaviridae in the new order, Bunyavirales. Most arboviruses possess an RNA genome and are mainly transmitted by mosquitoes (3). While acute infections in vertebrate hosts are typically self-limiting, arboviruses establish persistent infections in arthropods granting to the vector a central role as a viral reservoir (4).

Arboviruses circulate primarily within an enzootic cycle involving zoophilic vector species and non-human hosts. Recurrent spillovers cause occasional infections of humans initiating an epidemic cycle. Arboviruses such as dengue (DENV; *Flavivirus*, Flaviviridae), chikungunya (CHIKV; *Alphavirus*, Togaviridae), Zika (ZIKV; *Flavivirus*, Flaviviridae) and, Yellow fever virus (YFV; *Flavivirus*, Flaviviridae) do not need to amplify in wild animals to cause outbreaks in humans, which act simultaneously as amplifier, disseminator and source of infection for the major vectors, the anthropophilic mosquitoes *Aedes aegypti* and *Aedes albopictus* (5). Thus, the success of these viruses comes from their feature to be mainly transmitted by human-biting mosquitoes strongly adapted to urban environments. The establishment of a new epidemic cycle is undoubtedly related to the introduction of a viremic vertebrate host (humans, animals) acting as a vehicle for importation of the virus into environments receptive to viral amplification.

Other arboviruses such as West Nile virus (WNV; *Flavivirus*, Flaviviridae) remain circulating within an enzootic cycle with sporadic spillovers causing human cases.

Many regions experience simultaneous circulation of different arboviruses (6, 7), and co-infections in vectors were reported (8). These coinfections can present an opportunity for viruses to exchange genetic material. Impacts of such genetic events on virulence for vertebrate hosts are still unknown (9). Thus, being able to detect a wide range of arboviruses in thousands of field-collected mosquitoes in a single experiment can be a valuable tool to predict arboviral emergences in human populations. Indeed similar methods were developed with success to screen tick-borne pathogens (bacteria, parasites and viruses) and allowed the detection of expected and unexpected pathogens in large scale epidemiological studies (10, 11). Therefore, we developed a high throughput system based on real-time microfluidic PCR which is able to detect 96 mosquito-borne viruses in 96 samples within one single run. With this method, we have screened: (1) mosquitoes infected artificially using a feeding system to validate our tool, (2) mosquitoes collected in countries endemic for the major human arboviruses (e.g., Senegal, Cambodia, Brazil), and (3) mosquitoes collected during the Zika and Yellow fever outbreaks in the Americas (French Guiana, Guadeloupe, Brazil, Suriname). This method allowed detecting epidemic viruses (ZIKV, CHIKV, YFV) but also unexpected viruses (e.g. Trivittatus virus, TVTV, *Orthobunyavirus*, Bunyaviridae) underlining the need of such a tool for early detection of emerging mosquito-borne viruses.

## 2. Materials and methods

### 2.1. Mosquitoes

To test the ability of our assays to detect viruses present in pools of mosquitoes, 47 batches of three infected mosquitoes of the species, *Ae. aegypti* and *Ae. albopictus* (infection performed by artificial feeding system), were provided by the Institut Pasteur (Paris). Six different viruses, single or double infections, were tested in a pilot study.

In ZIKV-endemic and -epidemic regions from South America, Africa, and Asia (Brazil, French Guiana, Guadeloupe, Suriname, Senegal, Cambodia), adult mosquitoes were collected, identified using morphological characters and dissected to separate abdomen from the remaining body parts (RBP) (See Tables 1-6 for details). Abdomens of the same species were grouped by pools of 20-30 individuals in cryovials, and RBP were stored individually at −80°C until further analysis.

**Table 1.** Mosquito and sandflies species, number of mosquitoes collected and number of pools analyzed in Senegal.

| Stage collected | Collection Site | GPS coordinates | Urban<br>Rural<br>Sylvatic | Mosquito species | Number of arthropods screened | Virus detected through microfluidic system | Type of confirmation performed |
| --- | --- | --- | --- | --- | --- | --- | --- |
| Adult | Baraboye | 12°41'11.2"N/12°24'39.2"W | Rural | *Ae. furcifer*, *Ae. dalzieli*, *Ae. taylori*, *Ae. vittatus*,<br>*An. coustani*, *An. funestus*, *Ma. uniformis*, sandflies | 97 | - | - |
|  | Ngari | 12°38'07.3"N/12°14'59.5"W | Rural | *Ae. furcifer*, *Ae. dalzieli*, *Ae. aegypti*, *Ae. argenteopunctatus*, *Ae. hirsutus*, *Ae. mcintoshi*, *Cx. quinquefasciatus*, *Ae. vittatus*,<br>*An. coustani*, *An. funestus*, | 96 | - | - |
|  | Silling | 12°32'36.5"N/12°16'18.7"W | Rural | *Ae. luteocephalus*, *An. gambiae* | 3 | - | - |
|  | Tenkoto | 12°40'23.1"N/12°16'37.1"W | Rural | *Ae. furcifer* | 5 | - | - |
|  | Velingara | 12°27'33.9"N/12°03'17.3"W | Rural | *Ae. aegypti*, *Ae. dalzieli*, *Ae. furcifer*, *Ae. luteocephalus*,<br>*Ae. vittatus*, *An. coustani* | 43 | - | - |
|  | Kedougou (C1F) | 12°39'42.1"N/12°16'05.2"W | Sylvatic | *Ae. aegypti*, *Ae. furcifer*, *Ae. luteocephalus*, *Ae. taylori*, *Ae. unilineatus*, *Ae. vittatus*, *An. coustani*, *An. funestus*,<br>*An. nili*, *Cx. perfuscus*, *Ma. africana*, *Ma. uniformis* | 151 | - | - |
|  | Kedougou (D1F) | 12°36'43.9"N/12°14'50.7"W | Sylvatic | *Ae. aegypti*, *Ae. africanus*, *Ae. dalzieli*, *Ae. furcifer*, *Ae. luteocephalus*, *Ae. taylori*, *Ae. unilineatus*, *Ae. vittatus*, *An. coustani*, *An. funestus*, *An. nili*, *Cx. annulioris*, *Cx. bitaeniorhynchus*, *Cx. poicilipes*, *Cx. perfuscus*,<br>*Ma. africana*, *Ma. uniformis* | 278 | YFV* | YFV confirmed by PCR |
|  | Kedougou (E2F) | 12°29'21.2"N/12°06'06.2"W | Sylvatic | *Ae. aegypti*, *Ae. africanus*, *Ae. argenteopunctatus*, *Ae. dalzieli*, *Ae. furcifer*,<br>*Ae. luteocephalus*, *Ae. taylori*, *Ae. unilineatus*, *Ae. vittatus*, *An. coustani*, *An. funestus*,<br>*Cx. poicilipes*, *Ma. uniformis* | 261 | - | - |
|  | 8 |  |  |  | **934** |  |  |
| Larvae | Dalaba | 12°33'25.6"N/12°10'41.0"W | Rural | *Ae. aegypti* | 24 | - | - |
|  | Ngari | 12°38'07.3"N/12°14'59.5"W | Rural | *Ae. aegypti*, *Ae. vittatus* | 18 | - | - |
|  | Kedougou (D1F) | 12°36'43.9"N/12°14'50.7"W | Sylvatic | *Ae. aegypti*, *Ae. bromeliae*, *Ae. furcifer*, *Ae. longipalpis*,<br>*Ae. luteocephalus*, *Ae. taylori*, *Ae. unilineatus*,<br>*Ae. vittatus* | 237 | - | - |
|  | Kedougou (E2F) | 12°29'21.2"N/12°06'06.2"W | Sylvatic | *Ae. aegypti*, *Ae. bromeliae*, *Ae. longipalpis*,<br>*Ae. luteocephalus*, *Ae. neoaffricanus*, *Ae. taylori*,<br>*Ae. unilineatus*, *Ae. vittatus*, *Er. chrysogaster* | 123 | - | - |
|  | 4 |  |  |  | **402** |  |  |

*Ae.*, *Aedes*; *An.*, *Anopheles*; *Cx.*, *Culex*; *Er.*, *Eretmapodites*; *Ma.*, *Mansonia*; *YFV*, Yellow fever virus.
\* YFV detected in one pool of *Ae. scapularis*, one pool of *Ae. furcifer* and confirmed in one female of this pool.

### 2.2. RNA extraction

Total RNAs were extracted from each pool using the Nucleospin RNA II extraction kit (Macherey-Nagel, Germany). Pools were ground in 350 µL Lysis Buffer and 3.5 µL β-mercaptoethanol using the homogenizer Precellys®24 Dual (Bertin, France) at 5,500 rpm for 20 sec. Total RNA per pool was eluted in 50 µL of RNase free water and stored at −80°C until use.

When pools of abdomens were positive for virus, the RBP (head/thorax) of individual mosquitoes composing each pool were homogenized in 300 µL of DMEM with 10% fetal calf serum using the homogenizer Precellys®24 Dual (Bertin, France) at 5,500 rpm for 20 sec. Then total RNAs were extracted from 100 µL of homogenates using the Nucleospin RNA II extract kit (Macherey-Nagel, Germany) and 200 µL were conserved at −80°C for attempts to isolate the virus. Total RNA per sample was eluted in 50 µL of RNase free water and stored at −80°C until use.

### 2.3. Reverse Transcription and cDNA pre-amplification

RNAs were transcribed to cDNA by reverse transcription using the qScript cDNA Supermix kit according to the manufacturer’s instructions (Quanta Biosciences, Beverly, USA). Briefly, the reaction was performed in a final volume of 5 µL containing 1 µL of qScript cDNA supermix 5X, 1 µL of RNA and 3 µL of RNase free water; with one cycle at 25°C for 5 min, one cycle at 42°C for 30 min and one final cycle at 85°C for 5 min.

For cDNA pre-amplification, the Perfecta Preamp Supermix (Quanta Biosciences, Beverly, USA) was used according to the manufacturer’s instructions. All primers were pooled to a final concentration of 200 nM each. The reaction was performed in a final volume of 5 μL containing 1 μL Perfecta Preamp 5X, 1.25 μL pooled primers, 1.5 µL distilled water and 1.25 μL cDNA, with one cycle at 95°C for 2 min, 14 cycles at 95°C for 10 sec and 3 min at 60°C. At the end of the cycling program, the reactions were 1:5 diluted. Pre-amplified cDNAs were stored at −20°C until use.

### 2.4. Assay design

Mosquito-borne viruses (MBV), their targeted genes and the corresponding primers/probe sets are listed in Table 7. For a total of 64 viruses including 149 genotypes/serotypes, primers and probes were specifically designed. Each primer/probe set was validated using a dilution range of several cDNA positive controls (when available) (Table 7), by real-time PCR on a LightCycler® 480 (LC480) (Roche Applied Science, Germany). Real-time PCR assays were performed in a final volume of 12 µL using the LightCycler® 480 Probe Master Mix 1X (Roche Applied Science, Germany) with primers and probes at 200 nM and 2 µL of control cDNA (virus reference material) or DNA (Plasmid). Thermal cycling conditions were as follows: 95°C for 5 min, 45 cycles at 95°C for 10 sec and 60°C for 15 sec and one final cooling cycle at 40°C for 10 sec.

### 2.5. High-throughput real-time PCR

The BioMark^TM^ real-time PCR system (Fluidigm, USA) was used for high-throughput microfluidic real-time PCR amplification using the 96.96 dynamic arrays (Fluidigm, USA). These chips dispensed 96 PCR mixes and 96 samples into individual wells, after which on-chip microfluidics assemble PCR reactions in individual chambers prior to thermal cycling resulting in 9,216 individual reactions. Real-time PCRs were performed using FAM- and black hole quencher (BHQ1)-labeled TaqMan probes with TaqMan Gene Expression Master Mix in accordance with manufacturer’s instructions (Applied Biosystems, France). Thermal cycling conditions were as follows: 2 min at 50°C, 10 min at 95°C, followed by 40 cycles of 2-step amplification of 15 sec at 95°C, and 1 min at 60°C. Data were acquired on the BioMark^TM^ real-time PCR system and analyzed using the Fluidigm real-time PCR Analysis software to obtain C_t_ values (see Michelet et al. 2014 for more details (12)). Primers and probes were evaluated for their specificity against cDNA reference samples. One negative water control was included per chip. To determine if factors present in the sample could inhibit the PCR, *Escherichia coli* strain EDL933 DNA was added to each sample as an internal inhibition control, using primers and probe specific for the *E. coli eae* gene (13).

### 2.7 Validation of the results by real-time PCR, virus isolation and genome sequencing

When cDNA of pools of abdomens were detected positive for viruses, the cDNAs of RBP (head/thorax) of individual mosquitoes composing each pool were screened by real-time PCRs on a LightCycler® 480 (LC480) (Roche Applied Science, Germany). Real-time PCR assay targeting the virus of interest (see primers/probe sets in Table 7) was performed in a final volume of 12 µL using the LightCycler® 480 Probe Master Mix 1X (Roche Applied Science, Germany), with primers and probes at 200 nM and 2 µL of control DNA. Thermal cycling conditions were as follows: 95°C for 5 min, 45 cycles at 95°C for 10 sec and 60°C for 15 sec and one final cooling cycle at 40°C for 10 sec.

When a positive sample was confirmed, virus isolation was attempted in Vero and C6/36 cells. Then, total RNA was extracted using the Nucleospin RNA II extract kit (Macherey-Nagel, Germany) following the manufacturer instructions and full genome sequencing was attempted. For ZIKV, twelve overlapping amplicons were produced using the reverse transcriptase Platinum Taq High Fidelity polymerase enzyme (Thermo Fisher Scientific) and specific primers (Table 8). PCR products were pooled in equimolar proportions. After Qubit quantification using Qubit® dsDNA HS Assay Kit and Qubit 2.0 fluorometer (ThermoFisher Scientific) amplicons were fragmented (sonication) into fragments of 200 bp long. Libraries were built adding barcode for sample identification, and primers to fragmented DNA using AB Library Builder System (ThermoFisher Scientific). To pool the barcoded samples equimolarly, a quantification step by the 2100 Bioanalyzer instrument (Agilent Technologies) was performed. An emulsion PCR of the pools and loading on 520 chip was done using the automated Ion Chef instrument (ThermoFisher Scientific). Sequencing was performed using the S5 Ion torrent technology (Thermo Fisher Scientific) following manufacturer’s instructions. Consensus sequence was obtained after removing the thirty first and last nucleotides of each read, trimming reads depending on quality (reads with quality over >99%) and length (reads over 100 pb were kept) and mapping them on a reference (KY415987, most similar sequence after Blastn) using CLC genomics workbench software 11.0.1 (Qiagen). A *de novo* contig was also produced to ensure that the consensus sequence was not affected by the reference sequence.

## 3. Results

One hundred and forty-nine primer/probe sets were designed to detect 64 MBV (Table 7). Among them, 95 sets of primers/probe specifically identified their corresponding positive control samples (37 viral RNA) *via* Taqman RT-real-time PCRs or Taqman real-time PCRs on a LightCycler 480 apparatus. Resulting C_t_ values varied from 8 to 42 depending on sample type and nucleic acid concentration. Unfortunately, 54 designs were not tested due to the lack of RNA positive control.

To avoid sensitivity problems, cDNA pre-amplification was included in the assay. This step enabled detection of all positive controls (95 primer/probe sets tested on 37 viral RNAs) via Taqman real-time PCRs on a LC480 apparatus. The specificity of each primers/probe set was then evaluated using 37 MBV positive controls on the BioMark^TM^ system (Fig. 1). Results demonstrated high specificity for each primer/probe set after pre-amplification (Fig. 1). Indeed, 91 assays (among the 149 developed) were only positive for their corresponding positive controls. Four designs demonstrated cross-reactivity with a virus from the same species or genus: DENV-1 assay amplified also DENV-2, DENV-2 assay cross-reacted with DENV-3 and DENV-4, DENV-4 cross-reacted with DENV-3, one WNV assay amplified Usutu virus (USUV). Specificity of 54 assays was not fully tested in the absence of their respective positive controls. Nevertheless, those designs did not show any cross-reaction with RNA positive controls from other viruses.

**Figure 1.**
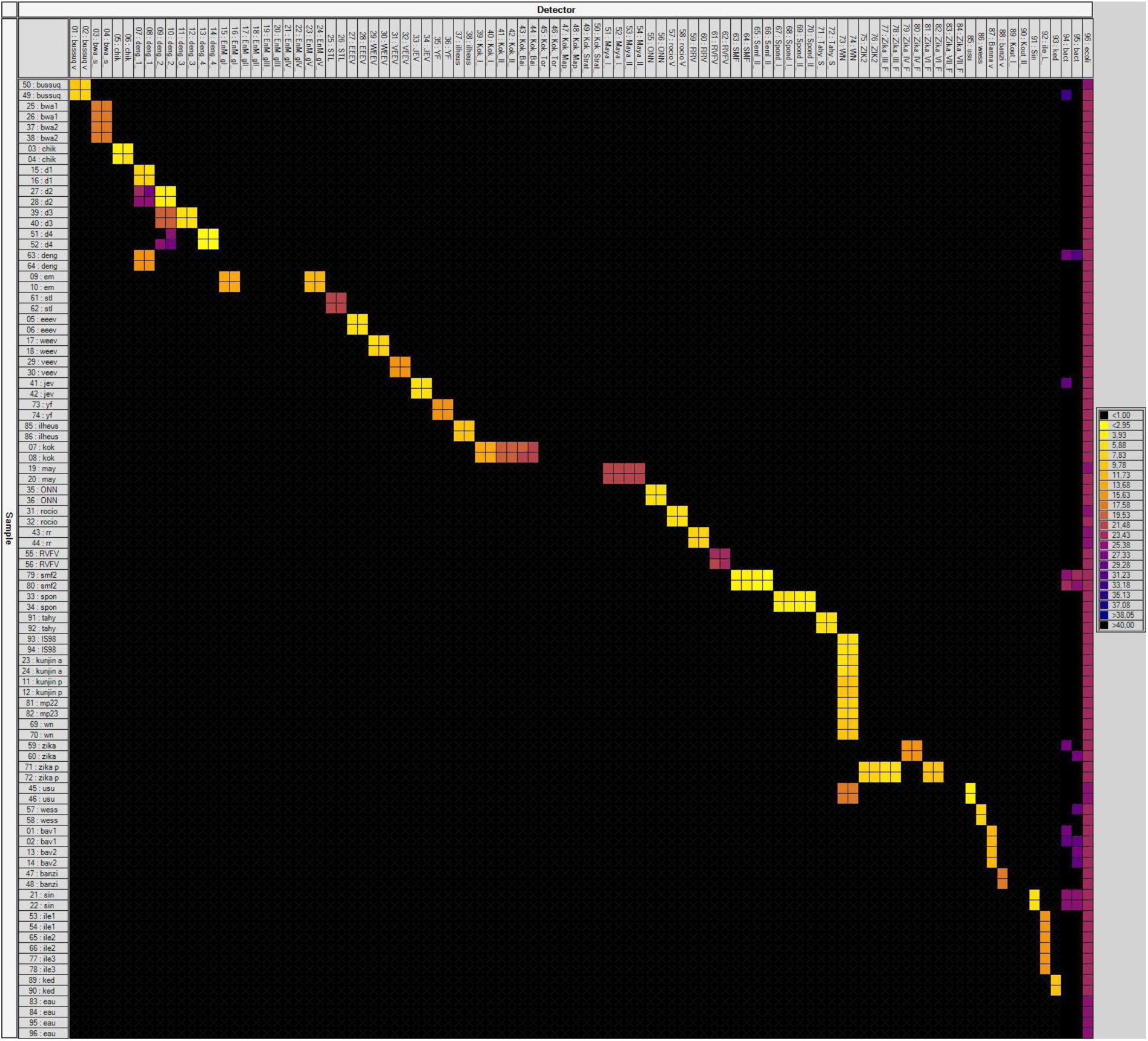
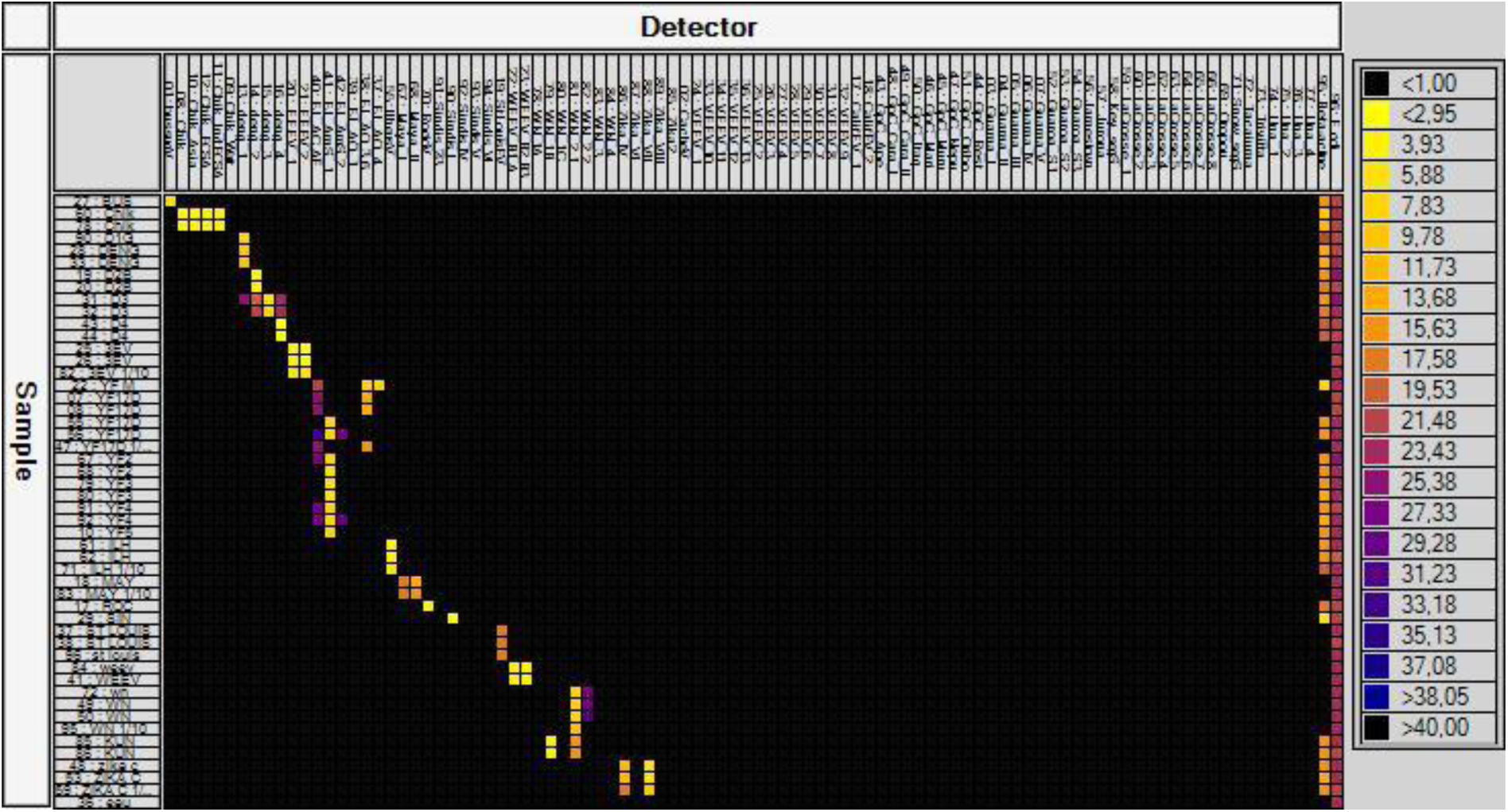
BioMark^TM^ dynamic array system specificity test (96.96 chip). Specificity of primers/probe sets from the Table 7 are presented into two figures 1A and 1B. Each square corresponds to a single real-time PCR reaction, where rows indicate the pathogen in the positive control and columns represent the targets of each primer/probe set. C_t_ values for each reaction are indicated in color; the corresponding color scale is presented in the legend on the right. The darkest shade of blue and black squares are considered as negative reactions with C_t_ > 30.

### Laboratory-infected mosquitoes

Forty-seven batches containing each, three infected mosquitoes, were screened with the high-throughput technic developed. The system was able to identify the six viruses present in different mosquitoes (Fig. 2). Indeed, seven batches were infected by DENV-1, four by DENV-3, four by DENV-4, 3 by CHIKV, five by WNV, 13 batches by ZIKV and 10 batches were coinfected by CHIKV and DENV-2. As for the specificity test, DENV-1, DENV-2 and DENV-3 assays demonstrated cross-reactions.

**Figure 2.**
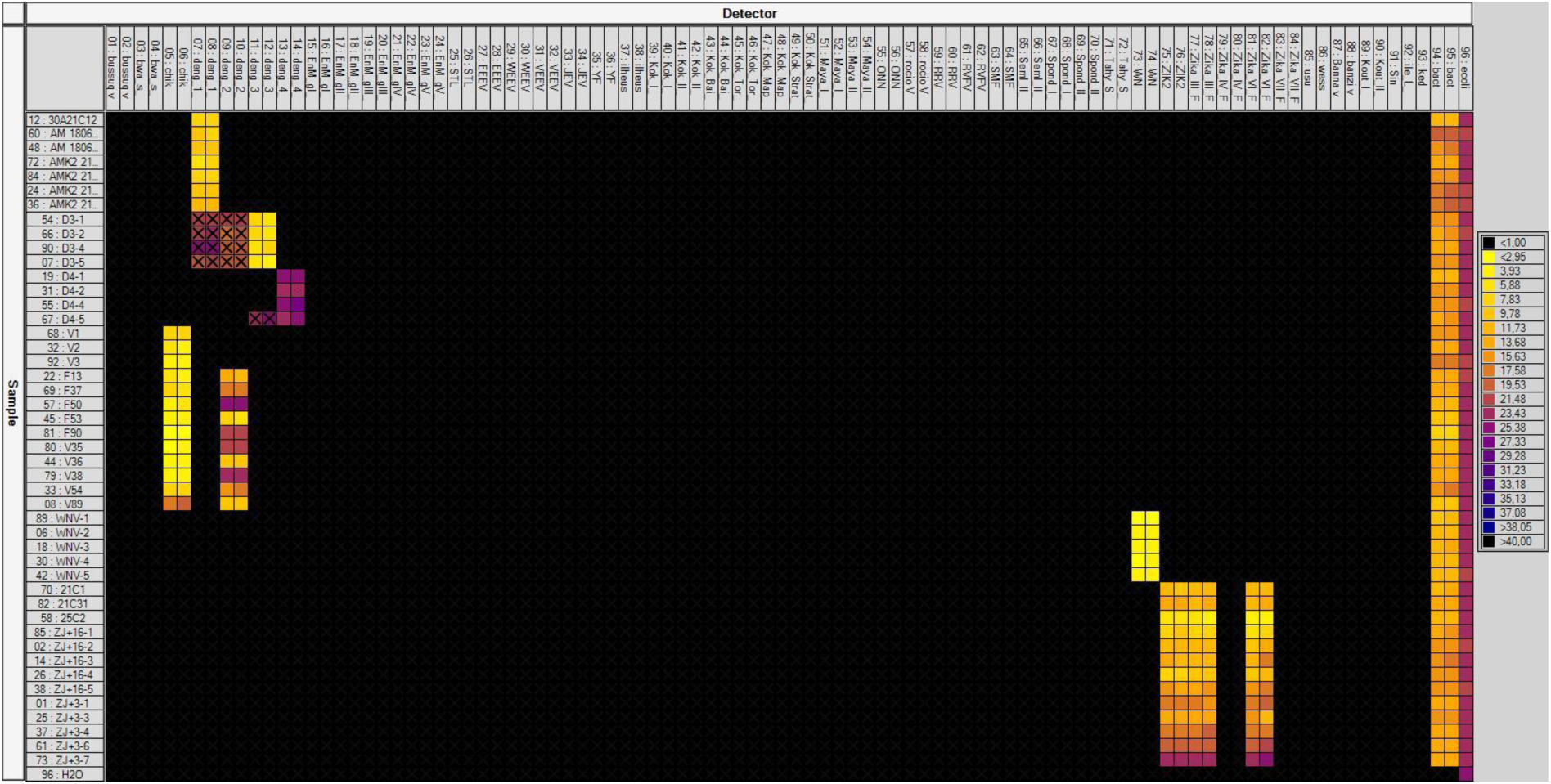
Screening of artificially infected mosquitoes through the BioMark^TM^ dynamic array system developed (96.96 chip). Each square corresponds to a single real-time PCR reaction, where rows indicate the batches of mosquitoes tested and columns represent the targets of each primer/probe set. Cross indicate cross-reaction of assays. C_t_ values for each reaction are indicated in color; the corresponding color scale is presented in the legend on the right. The darkest shade of blue and black squares are considered as negative reactions with C_t_ > 30.

### Field-collected mosquitoes from endemic and epidemic areas

A total of 17,958 field-collected mosquitoes in six countries from the African, American and Asian continents were screened for arbovirus.

### Endemic areas

#### Senegal

In Senegal, 934 arthropods including 6 sandflies and 928 mosquitoes (25 males and 909 females) from 21 species and five genera (detailed in Table 1), were collected in the Kedougou area (Southeastern Senegal) from August to November 2017. Moreover, 402 larvae were also collected in the same area from August 2017 to January 2018 and reared until adult emergence in insectarium (188 males and 214 females obtained). Mosquitoes were grouped by species and sex; 231 and 112 pools were respectively analyzed for MBVs. YFV was detected in one pool of 20 females of *Aedes furcifer* and was confirmed in head/thorax from 1 *Aedes furcifer* female by RT-real-time PCR. Virus was identified as YFV from West Africa lineage currently circulating in Senegal (38). Virus isolation was attempted but without any success.

#### Cambodia

In Cambodia, 492 mosquitoes (73 males and 419 females) from 28 species and 5 genera (detailed in Table 2), were collected in one area at two periods, the dry season in May 2019 and the rainy season in November 2018. Mosquitoes were grouped by species and sex into 109 pools and were analyzed for MBVs. No virus was detected (Table 2).

**Table 2.** Mosquito species, number of mosquitoes collected and number of pools analyzed in Cambodia.

| Collection Site | GPS coordinates | Urban<br>Rural<br>Sylvatic | Mosquito species | Number of mosquitoes screened | Virus detected through microfluidic system | Type of confirmation performed |
| --- | --- | --- | --- | --- | --- | --- |
| Mondulkiri, Cambodia | 12°10'28.4" / 106°53'40.9" | Sylvatic | *Ae. aegypti*, *Ae. albopictus*, *Ae. gardnerri imitator*, *Ae. prominens*, *An. barbirostris*, *An. indefinitus*, *An. jamesii*, *An. kochi*, *An. mimulus complex*, *An. philippinensis*, *An. roperi*, *An. umbrosus*, *An. vegus*, *Ar. annulipalpis*, *Ar. dolichocephalus*, *Ar. flavus*, *Ar. foliatus/kuchingensis*, *Ar. moultoni*, *Ar. subalbatus*, *Ar. theobaldi*, *Cx. bitaeniorhynchus*, *Cx. brevipalpis*, *Cx. fuscocephala*, *Cx. perplexus/whitei*, *Cx. sitiens*, *Cx. vishnui complex*, *Hz. catesi*, *Hz. demeilloni* | 492 | - | - |

*Ae.*, *Aedes*; *An.*, *Anopheles*; *Ar.*, *Armigeres*; *Cx.*, *Culex*; *Hz.*, *Heizmannia*.

### Endemic/epidemic area, Brazil

In Brazil, 7,705 mosquitoes (889 males and 6,816 females) belonging to 22 species and 15 genera (detailed in Table 3), were collected in 15 areas from January 2016 to May 2017. Mosquitoes were then grouped into 647 pools and were analyzed for MBVs. In total, three different viruses (YFV, CHIKV and Trivittatus virus (TVTV)) were preliminary detected in six pools (in 4, 1 and 1 respectively). Only the presence of YFV was confirmed in the head/thorax from individual mosquitoes, from three species (*Ae. scapularis, Ae. taeniorhynchus* and *Hg. leucocelaenus*) by RT real-time PCR corresponding to YFV strains currently circulating in Brazil (37). Attempts to isolate the virus were made but remained unsuccessful.

**Table 3.** Mosquito species, number of mosquitoes collected and viruses detected in Brazil.

| Collection Site | GPS coordinates | Urban<br>Rural<br>Sylvatic | Mosquito species | Number of<br>mosquitoes<br>screened | Virus detected<br>through<br>microfluidic<br>system | Type of confirmation<br>performed |
| --- | --- | --- | --- | --- | --- | --- |
| Belo Horizonte | 19°51'59.29"S/44° 0'43.51"W | Urban Forest | *Ae. albopictus*, *Ae. aegypti*,<br>*Cx. quinquefasciatus*, *Sa.*<br>*albiprivus* | 17 | - | - |
| Casimiro de Abreu | 22°26'33.31"S/42°12'30.34"W | Sylvatic | *Ae. scapularis* | 24 | - | - |
| Domingos Martins | 20°17'12.48"S/40°50'14.35"W | Sylvatic | *Ae. albopictus* | 7 | - | - |
| Goiânia | 6°40'16.32"S/49°22'49.93"W | Urban Forest | *Aedeomiya*, *Ae. aegypti*, *Ae.*<br>*albopictus*, *Aedes* sp.,<br>*Coquillettidia* sp., *Culex* sp.,<br>*Hg. leucocelaenus*, *Limatus*<br>sp., *Mansonia* sp.,<br>*Orthopodomyia* sp.,<br>*Psorophora* sp., *Sabethes* sp.,<br>*Wyeomyia* sp. | 689 | - | - |
| Guapimirim | 22°28'56.31"S/42°59'26.36"W | Sylvatic | *Ru. frontosa* | 10 | - | - |
| Macaé | 22°18'17.54"S/42° 0'8.80"W | Sylvatic | *Ae. scapularis*, *Wyeomyia* sp. | 25 | - | - |
| Manaus | 3°00'12.78"S/59°55'37.86"W | Urban Forest | *Ae. aegypti*, *Ae. albopictus*,<br>*Aedes* sp., *Culex* sp., *Hg.*<br>*leucocelaenus*, *Limatus* sp.,<br>*Orthopodomyia* sp.,<br>*Psorophora* sp., *Sabethes* sp.,<br>*Trichoposopum* sp.,<br>*Uranotenia* sp., *Wyeomyia* sp. | 3939 | CHIKV* | Isolation, conventional<br>and real-time PCR<br>failed to confirm the<br>result |
| Maricá | 22°55'24.44"S/42°42'27.88"W | Sylvatic | *Ae. aegypti*, *Ae. albopictus*,<br>*Ae. scapularis*, *Ae.*<br>*taeniorhynchus*, *Culex* sp.,<br>*Cx. nigripalpus*, *Hg.*<br>*janthinomys*, *Hg.*<br>*leucocelaenus*, *Li. durhamii*,<br>*Ru. humboldti* | 198 | YFV§ | YFV confirmed by<br>PCR |
| Miguel Pereira | 22°29'3.21"S/43°18'15.98"W | Sylvatic | *Sh. fluviatilis* | 12 | - | – |
| Nova Friburgo | 22°24'46.35"S/42°18'58.57"W | Sylvatic | *Ru. humboldti* | 2 | CHIKV<sup>£</sup> | Isolation, conventional and real-time PCR failed to confirm the result |
| Queluz | 22°41'52.33"S/44°43'43.91"W | Sylvatic | *Wy. pilicauda*, *Wy. confusa* | 6 | - | - |
| Rio de Janeiro | 22°52'45.7"S/43°18'10.0"W | Urban | *Ae. aegypti*, *Ae. albopictus*, *Cx. quinquefasciatus* | 261 | - | - |
| | 22°56'6.57"S/43°26'42.19"W | Urban Forest | *Ae. aegypti*, *Ae. albopictus*, *Aedes sp.*, *Coquillettidia sp.*, *Culex sp.*, *Hg. leucocelaenus*, *Mansonia sp.*, *Psorophora sp.*, *Runchomyia sp.*, *Sabethes sp.*, *Trichoposopum sp.*, *Wyeomyia sp.* | 2447 | TVTV<sup>\$</sup> | Isolation, conventional and real-time PCR failed to confirm the result |
| Serra | 20° 6'46.89"S/40°11'12.53"W | Sylvatic | *Ae. albopictus*, *Cx. quinquefasciatus* | 26 | - | - |
| Simonésia | 19°55'12.06"S/41°54'20.23"W | Sylvatic | *Ae. albopictus*, *Cq. venezuelensis*, *Hg. janthinomys*, *Hg. leucocelaenus*, *Sa. albiprivus* | 41 | - | - |
| Teresópolis | 22°26'58.56"S/42°59'5.43"W | Sylvatic | *Ru. frontosa* | 1 | - | - |
| **15** |  |  |  | **7705** |  |  |

*Ae.*, *Aedes*; *Cq.*, *Coquillettidia*; *Cx.*, *Culex*; *Hg.*, *Haemagogus*; *Li.*, *Limatus*; *Ru.*, *Runchomyia*; *Sa.*, *Sabethes*; *Sh.*, *Shannoniana*; *Wy.*, *Wyeomyia*; *CHIKV*, *Chikungunya virus*; *YFV*, *Yellow fever virus*; *TVTV*, *Trivittatus virus*. \*CHIKV detected in one pool of *Cx. erraticus*; <sup>\$</sup> YFV detected in one pool of *Ae. scapularis*, one pool of *Ae. taeniorhynchus*, one pool of *Hg. leucocelaenus*; <sup>£</sup> CHIKV detected in one pool of *Ru. humboldti*; <sup>\$</sup> TVTV detected in one pool of *Cx. nigripalpus*

### Epidemic areas

#### Guadeloupe

In Guadeloupe, 150 mosquito pools corresponding to 2,173 mosquitoes (884 males and 1,289 females), from five species (*Ae. aegypti*, *Culex quinquefasciatus*, *Anopheles albimanus*, *Cx. bisulcatus*, *Cx. nigripalpus*, 54 *Culex.* spp.) collected from May to June 2016 were screened for 64 MBVs. ZIKV was found in two pools of *Cx. quinquefasciatus* females and nine pools of *Ae. aegypti* (eight pools of females and one pool of males) (Table 4). ZIKV was detected only in the head/thorax of individual *Ae. aegypti* females from the eight positive pools by a RT-real-time PCR. Virus was isolated on Vero cells and full genome sequencing identified the Asian genotype (GenBank Accession Numbers: MN185324, MN185325, MN185327, MN185329, MN185330, MN185331, MN185332).

**Table 4.** Mosquito species, number of mosquitoes collected and virus detected in Guadeloupe.

| Collection Site | GPS coordinates | Urban<br>Rural<br>Sylvatic | Mosquito species | Number of mosquitoes screened | Virus detected through microfluidic system | Type of confirmation performed |
| --- | --- | --- | --- | --- | --- | --- |
| Gosier | 16° 12' 21.229"N/61° 29' 31.438"W | Urban | *Ae. aegypti*, *Cx. quinquefasciatus*,<br>*An. albimanus* | 399 | - | - |
| Deshaies | 16° 18' 24.973"N/61° 47' 39.556"W | Urban/Periurban | *Ae. aegypti*, *Cx. quinquefasciatus*,<br>*An. albimanus*, *Cx. bisulcatus*, *De. magnus* | 306 | ZIKV* | Confirmed by real-time PCR on head-thorax of individual mosquitoes, isolation of the virus and full genome sequencing |
| Petit Bourg | 16° 11' 29.476"N/61° 35' 25.753"W | Urban/Periurban | *Ae. aegypti*, *Cx. quinquefasciatus*,<br>*Cx. nigripalpus*, *Culex sp.* | 422 | ZIKV* | Confirmed by real-time PCR on head-thorax of individual mosquitoes, isolation of the virus and full genome sequencing |
| Le Moule | 16° 19' 52.342"N/61° 20' 37.41"W | Urban/Periurban | *Ae. aegypti*, *Cx. quinquefasciatus*,<br>*Culex sp.* | 202 | ZIKV* | Confirmed by real-time PCR on head-thorax of individual mosquitoes, isolation of the virus and full genome sequencing |
| Saint François | 16° 15' 5.141"N/61° 16' 26.825"W | Urban | *Ae. aegypti*, *Cx. quinquefasciatus* | 356 | ZIKV* | Confirmed by real-time PCR on head-thorax of individual mosquitoes, isolation of the virus and full genome sequencing |
| Sainte Anne | 16° 13' 31.613"N/61° 23' 9.377"W | Urban/Periurban | *Ae. aegypti*, *Cx. quinquefasciatus*,<br>*Cx. nigripalpus*, *Culex sp.* | 325 | ZIKV* | Confirmed by real-time PCR on head-thorax of individual mosquitoes, isolation of the virus and full genome sequencing |
| Baie Mahault | 16° 16' 3.979"N/61° 35' 13.337"W | Urban | *Ae. aegypti*, *Cx. quinquefasciatus* | 19 | - | - |
| Le Lamentin | 16° 16' 17.36"N/61° 37' 59.754"W | Urban/Periurban | *Ae. aegypti*, *Cx. quinquefasciatus* | 27 | - | - |
| Goyave | 16° 7' 26.447"N/61° 34' 40.253"W | Urban/Periurban | *Ae. aegypti*, *Cx. quinquefasciatus*,<br>*Culex sp.* | 86 | - | - |
| Morne-à-L'eau | 16° 19' 53.832" N/61° 27' 25.855"W | Urban | *Ae. aegypti*, *Cx. quinquefasciatus* | 19 | - | - |
| Pointe-à-Pître | 16° 14' 54.499"N/61° 32' 18.888"W | Urban | *Ae. aegypti*, *Cx. quinquefasciatus* | 5 | - | - |
| Saint-Claude | 16° 1' 36.077"N/61° 42' 6.703"W | Urban/Periurban | *Ae. aegypti* | 1 | - | - |
| Petit Canal | 16° 22' 49.03"N/61° 29' 14.384"W | Urban/Periurban | *Culex sp.* | 6 | - | - |
| **13** |  |  |  | **2173** |  |  |

*Ae.*, *Aedes*; *An.*, *Anopheles*; *Cx.*, *Culex*; *De.*, *Deinocerites*. \*ZIKV detected in 2 pools of *Cx. quinquefasciatus* and 9 pools of *Ae. aegypti*, and confirmed in 9 females *Ae. aegypti* (one female per pool).

#### French Guiana

In French Guiana, 3,942 mosquitoes (1,098 males and 2,844 females) from seven species (*Ae. aegypti, Ae. scapularis, Ae. taeniorhynchus*, *Cx. quinquefasciatus, Ma. titillans, Cq.venezualensis,* and *Cq. albicosta*) were collected in the Cayenne area from June to August 2016 and grouped into 248 pools to be screened. Three pools of *Ae. aegypti* and one pool of *Cx. quinquefasciatus* were detected positive for ZIKV (Table 5). After screening individual head/thorax from those pools, only pools of *Ae. aegypti* were confirmed positive. ZIKV was isolated and fully sequenced; it belonged to the Asian genotype (GenBank Accession numbers: MN185326 and MN185328).

**Table 5.** Mosquito species, number of mosquitoes collected and virus detected in French Guiana.

| Collection Site | GPS coordinates | Urban<br>Rural<br>Sylvatic | Mosquito species | Number of<br>mosquitoes<br>screened | Virus detected<br>through Microfluidic<br>system | Type of confirmation<br>performed |
| --- | --- | --- | --- | --- | --- | --- |
| Cayenne | 4°55'53.08"N/<br>52°18'55.99"W | Urban | *Ae. aegypti*, *Ae. scapularis*, *Cx. quinquefasciatus*, *Ma. titillans*, *Cq. venezualensis*, *Cq. albicosta* | 1928 | ZIKV* | Confirmed by real-time PCR on head-thorax of individual mosquitoes, isolation of the virus and full genome sequencing |
| Remire-Montjoly | 4°53'34.01"N/<br>52°16'34.32"W | Urban/Periurban | *Ae. aegypti*, *Ae. scapularis*, *Cx. quinquefasciatus*, *Ma. titillans*, *Cq. venezualensis* | 1078 | – | - |
| Matoury | 4°50'52.22"N/<br>52°19'41.58"W | Urban/Periurban | *Ae. aegypti*, *Ae. taeniorhynchus*, *Cx. quinquefasciatus*, *Ma. titillans*, *Cq. venezualensis*, *Cq. albicosta* | 936 | ZIKV* | Confirmed by real-time PCR on head-thorax of individual mosquitoes, isolation of the virus and full genome sequencing |
| 3 |  |  |  | 3942 |  |  |

*Ae.*, *Aedes*; *Cq.*, *Coquillettidia* ; *Cx.*, *Culex*; *Ma.*, *Mansonia*. \*ZIKV in 1 pools of *Cx. quinquefasciatus* and 3 pools of *Ae. aegypti*, and confirmed in 3 females *Ae.aegypti* (one female per pool)

#### Suriname

In Suriname, from March to May 2017, four species/genus of mosquitoes (2,256 *Ae. aegypti*, 29 *Culex* spp., 5 *Haemogogus* spp., 20 undetermined species) representing 2,310 adults, were grouped into 77 pools and screened. No virus was detected (Table 6).

**Table 6.** Mosquito species, number of mosquitoes collected and virus detected in Suriname.

| Collection Site | GPS | Urban<br>Rural<br>Sylvatic | Mosquito species | Number of<br>mosquitoes<br>screened | Virus detected<br>through<br>Microfluidic<br>system | Type of<br>confirmation<br>performed |
| --- | --- | --- | --- | --- | --- | --- |
| Paramaribo | 5°51'54.2"N/55°11'33.4"W | Urban (Paramaribo) | Undetermined | 4 | - | - |
| Roti shop | 5°51'59.616"N/55°6'20.952W | Urban (Paramaribo) | *Ae. aegypti* | 68 | - | - |
| Car mechanic | 5°50'35.8"N 55°06'56.7"W | Urban (Paramaribo) | *Ae. aegypti* | 96 | - | - |
| Kwikfit car<br>mechanic | 5°50'38.1"N 55°07'23.3"W | Urban (Paramaribo) | *Ae. aegypti* | 567 | - | - |
| Family home | 5°50'33.3"N 55°07'17.5"W | Urban (Paramaribo) | *Ae. aegypti* | 296 | - | - |
| Outpatient<br>clinic | 5°50'30.3"N/55°7'8.615"W | Urban (Paramaribo) | *Ae. aegypti* | 103 | - | - |
| Chi min<br>restaurant | 5°49'54.408"N/55°8'24.683W | Urban (Paramaribo) | *Ae. aegypti* | 78 | - | - |
| Albertine<br>retirement home | 5°48'49.572"N/55°11'27.6"W | Urban (Paramaribo) | *Ae. aegypti* | 661 | - | - |
| Medisch<br>Opvoedkundig<br>Bureau (MOB) | 5°49'43.212"N/55°10'41.375"W | Urban (Paramaribo) | *Ae. aegypti* | 376 | - | - |
| Brownsweg | 5°0'57.384"N/55°10'2.172"W | Rural/Sylvatic | *Ae. aegypti*, *Culex*<br>sp., Undetermined | 21 | - | - |
| Brownsberg | 4°56'36.24"N/55°10'6.6"W | Rural/Sylvatic | *Ae. aegypti*, *Culex*<br>sp., *Haemagogus*<br>sp., Undetermined | 40 | - | - |
| **11** |  |  |  | **2310** |  |  |

*Ae.*, *Aedes*; *Cx.*, *Culex*; *Hg.*, *Haemagogus*.

**Table 7.**
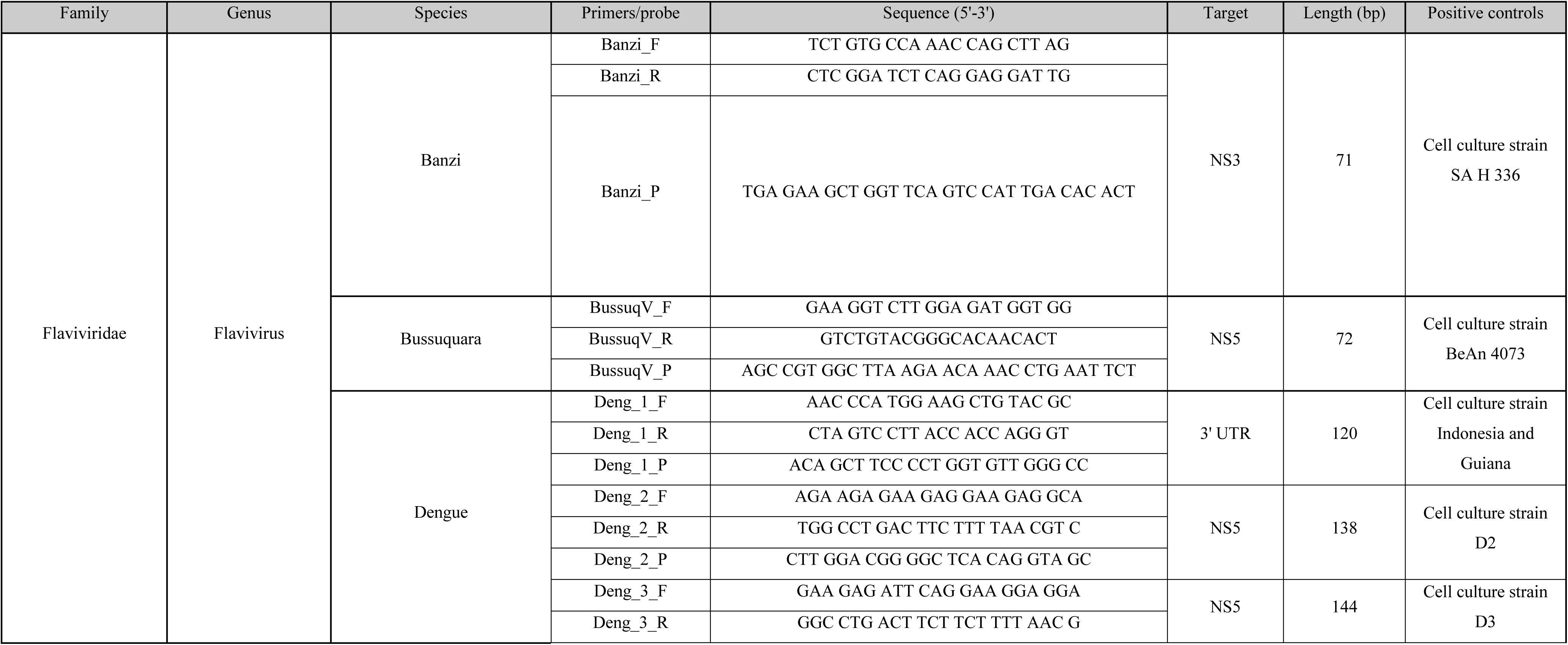
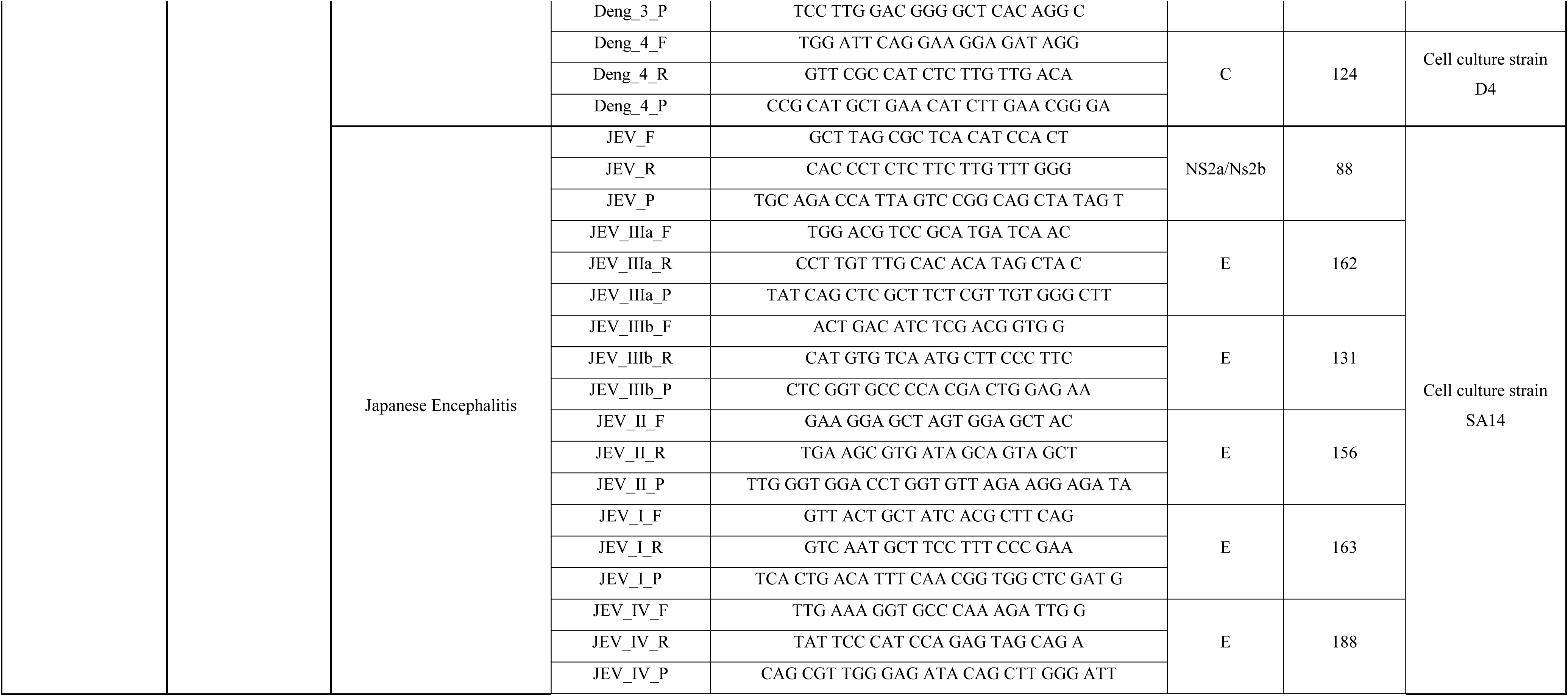
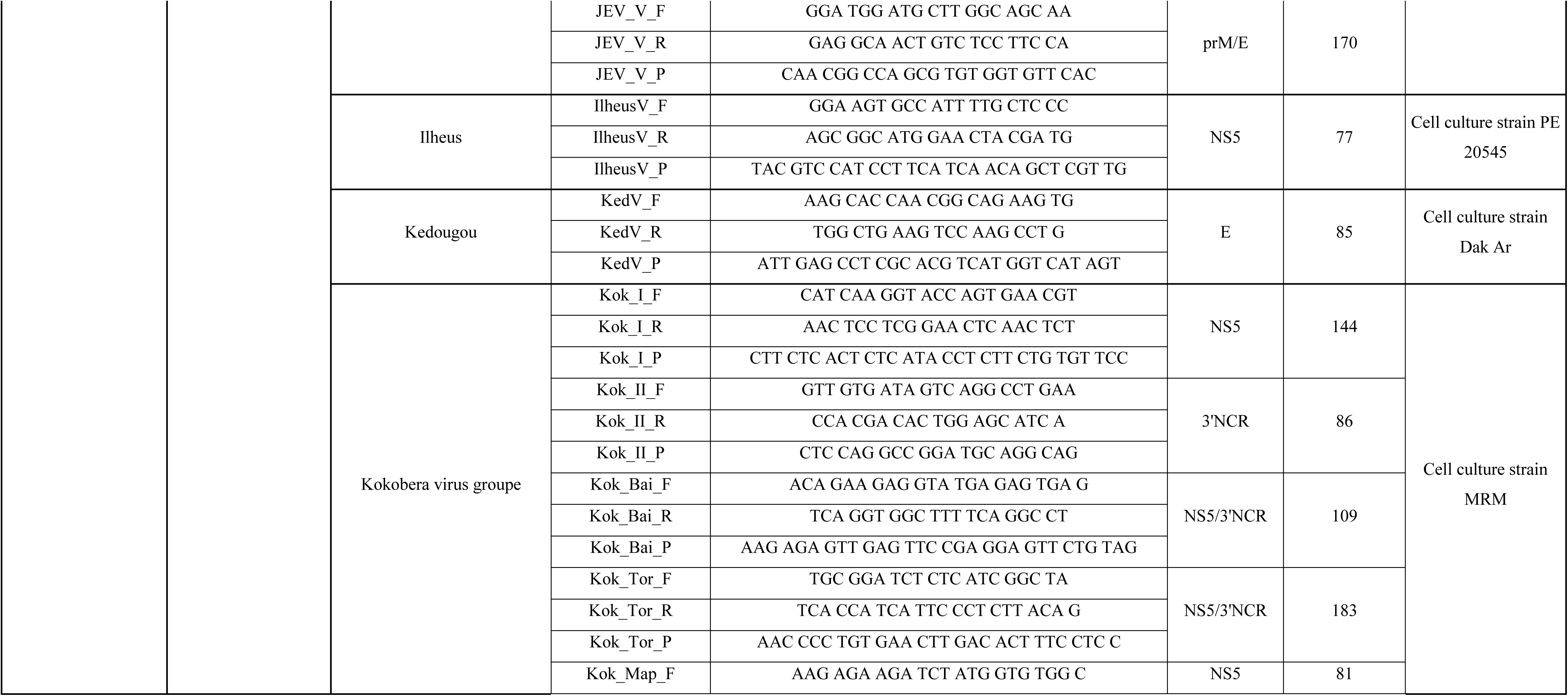
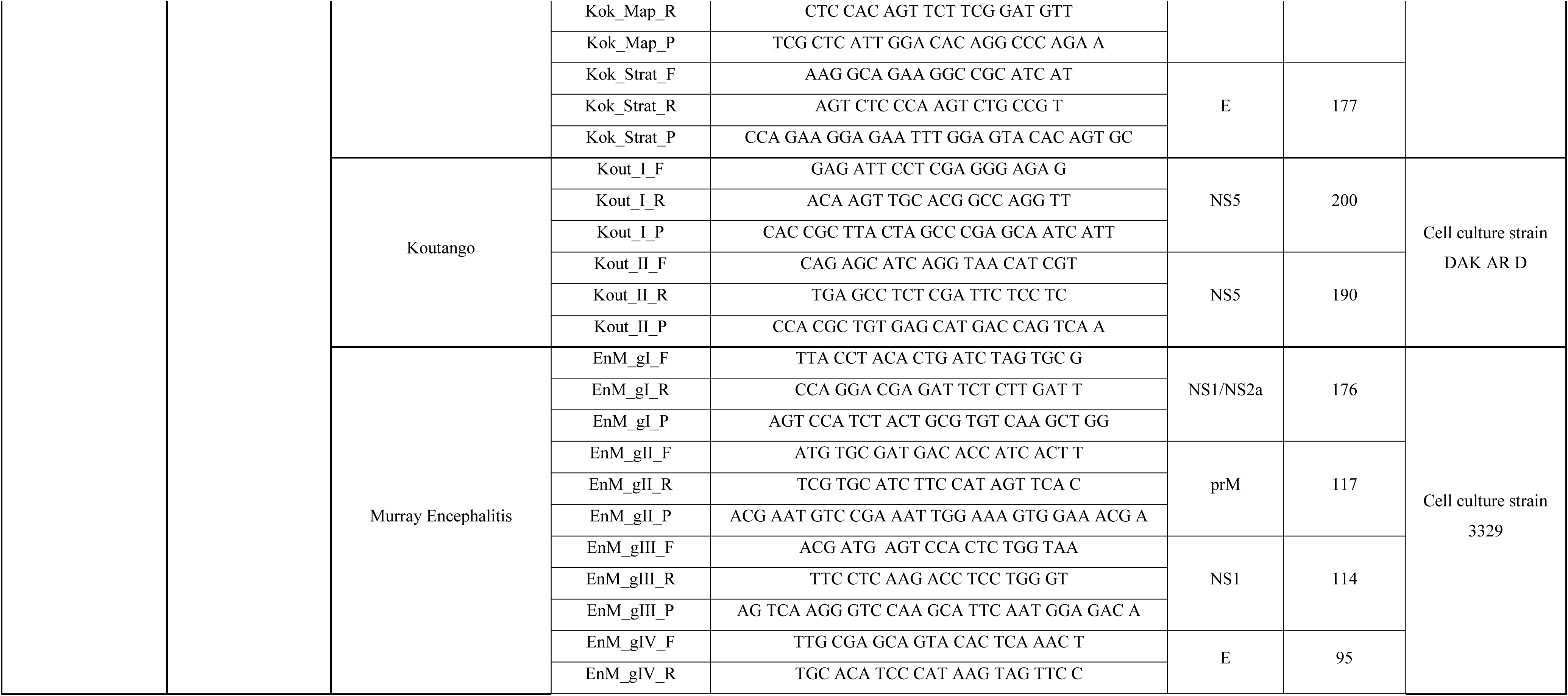
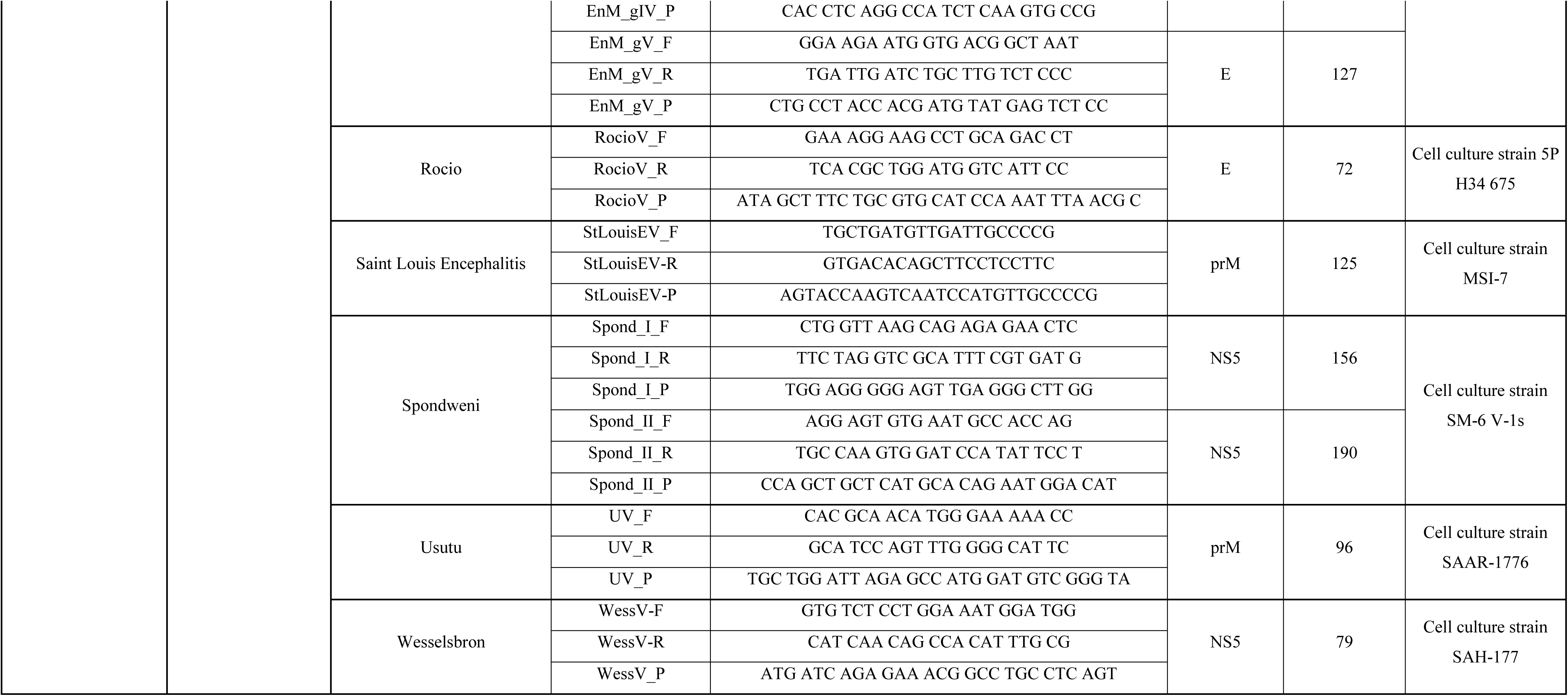
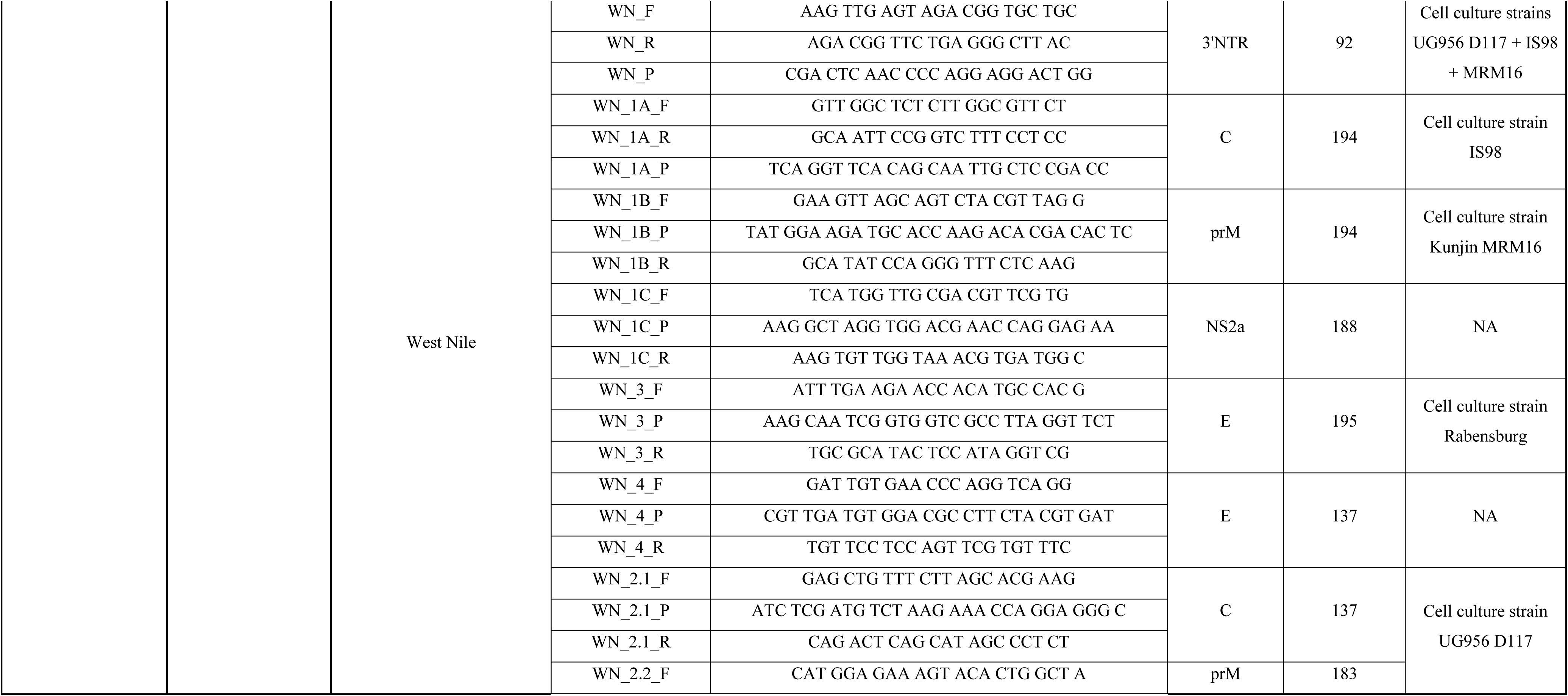
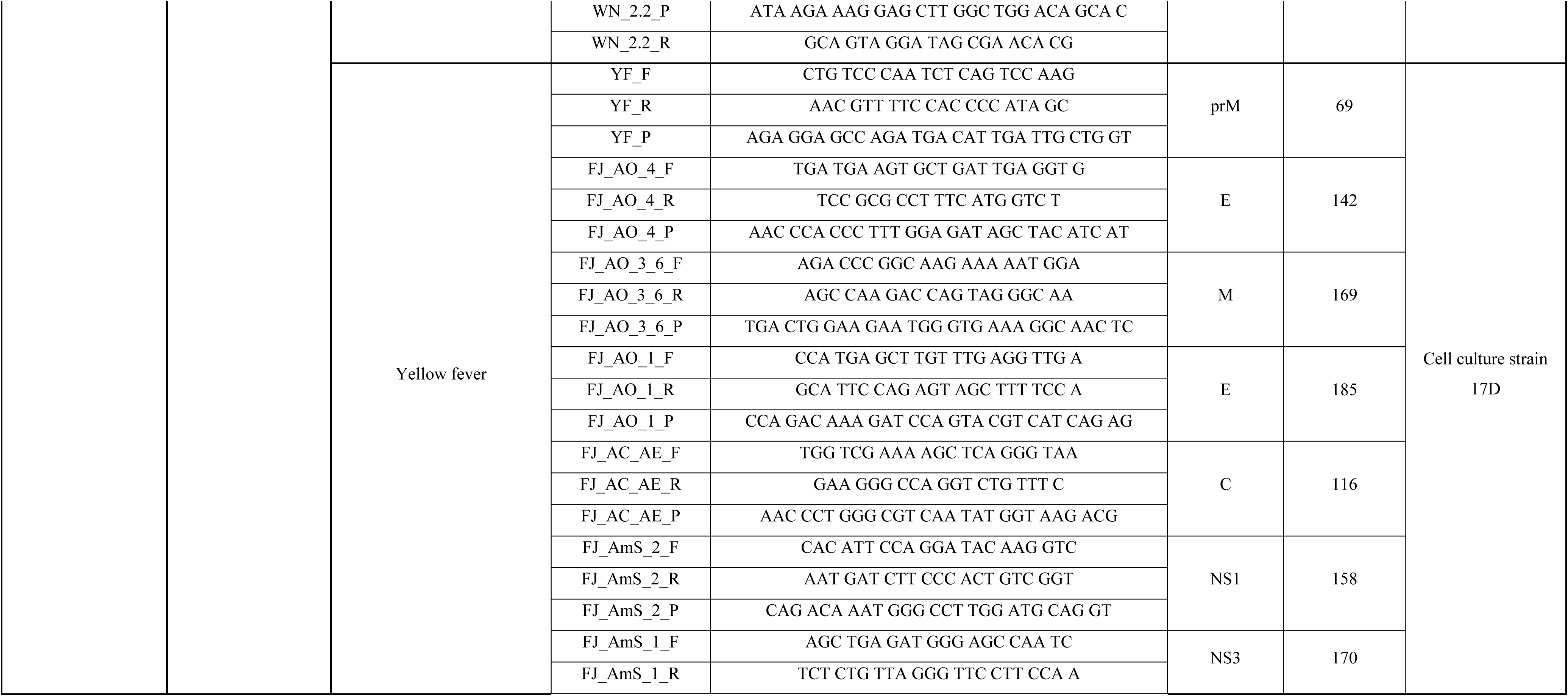
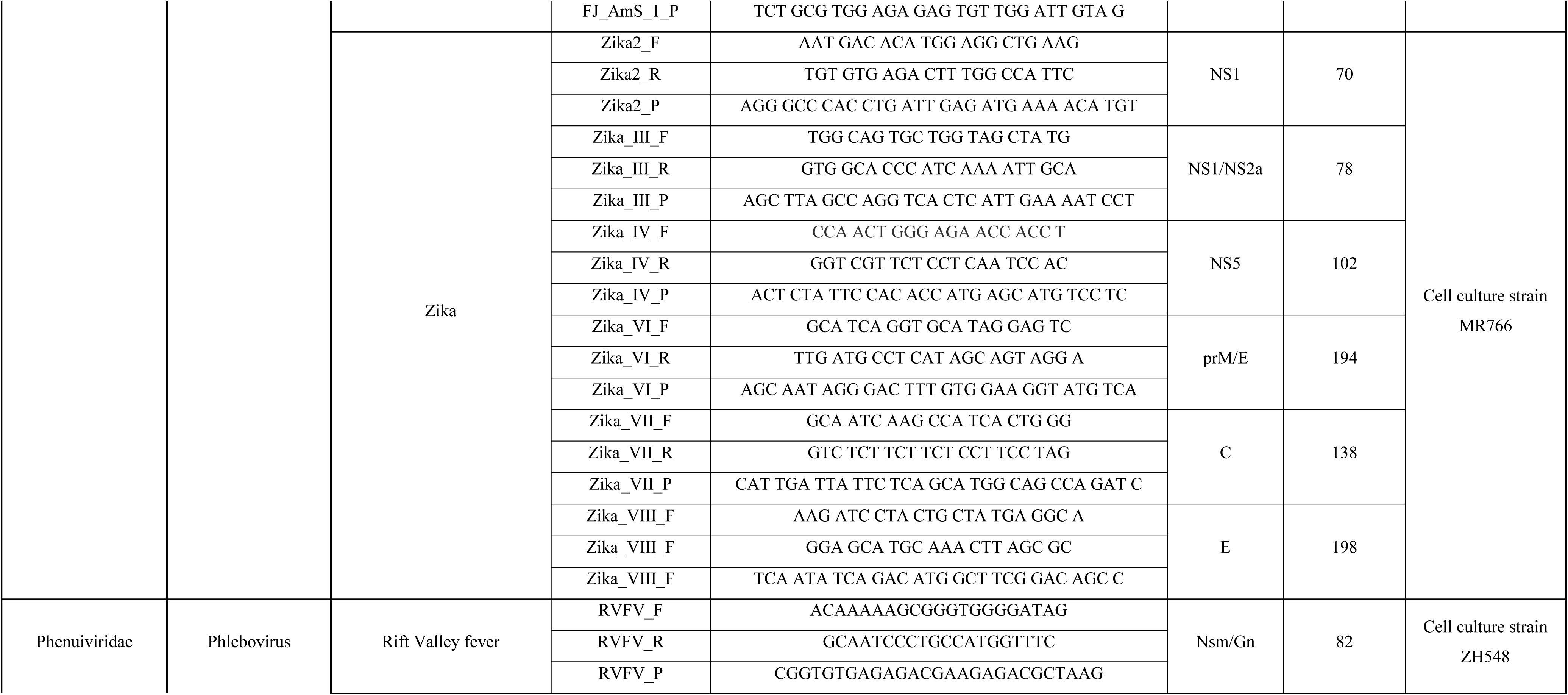
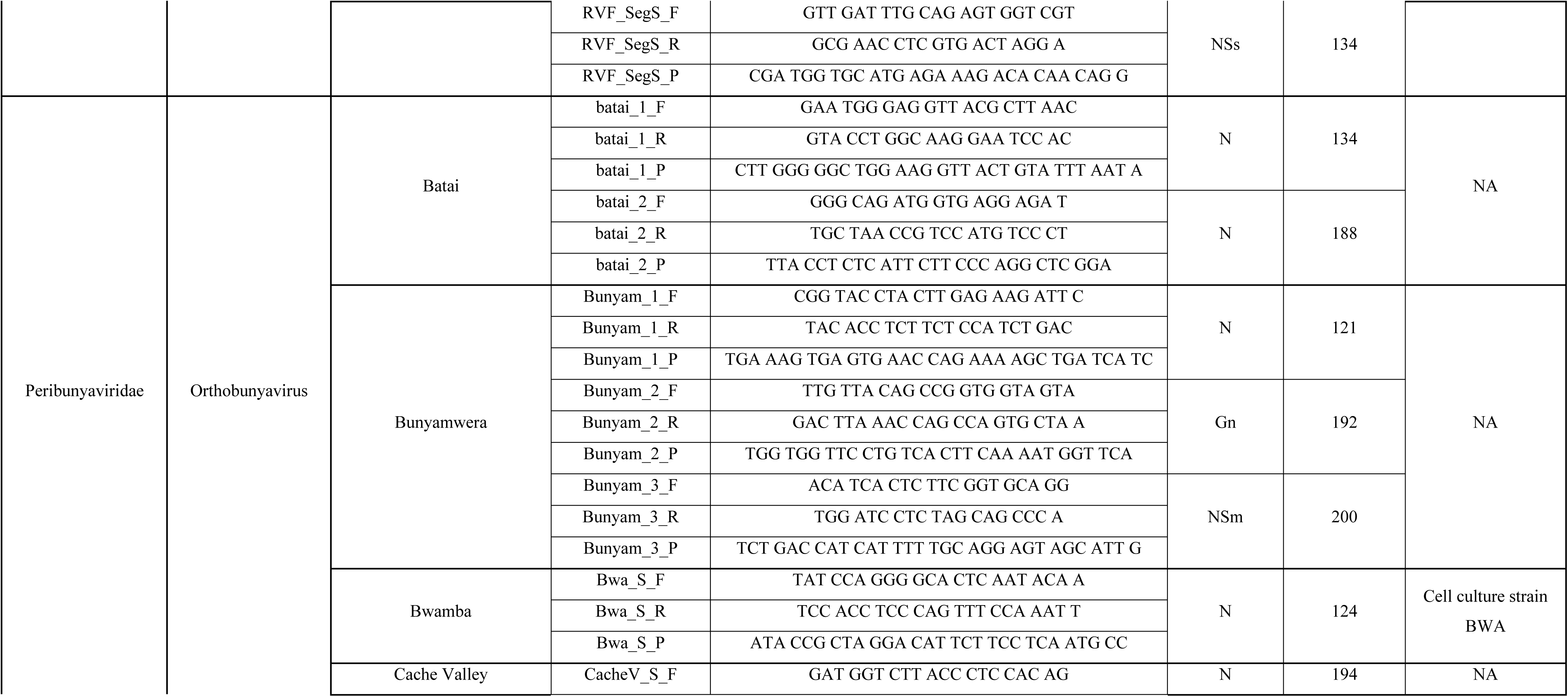
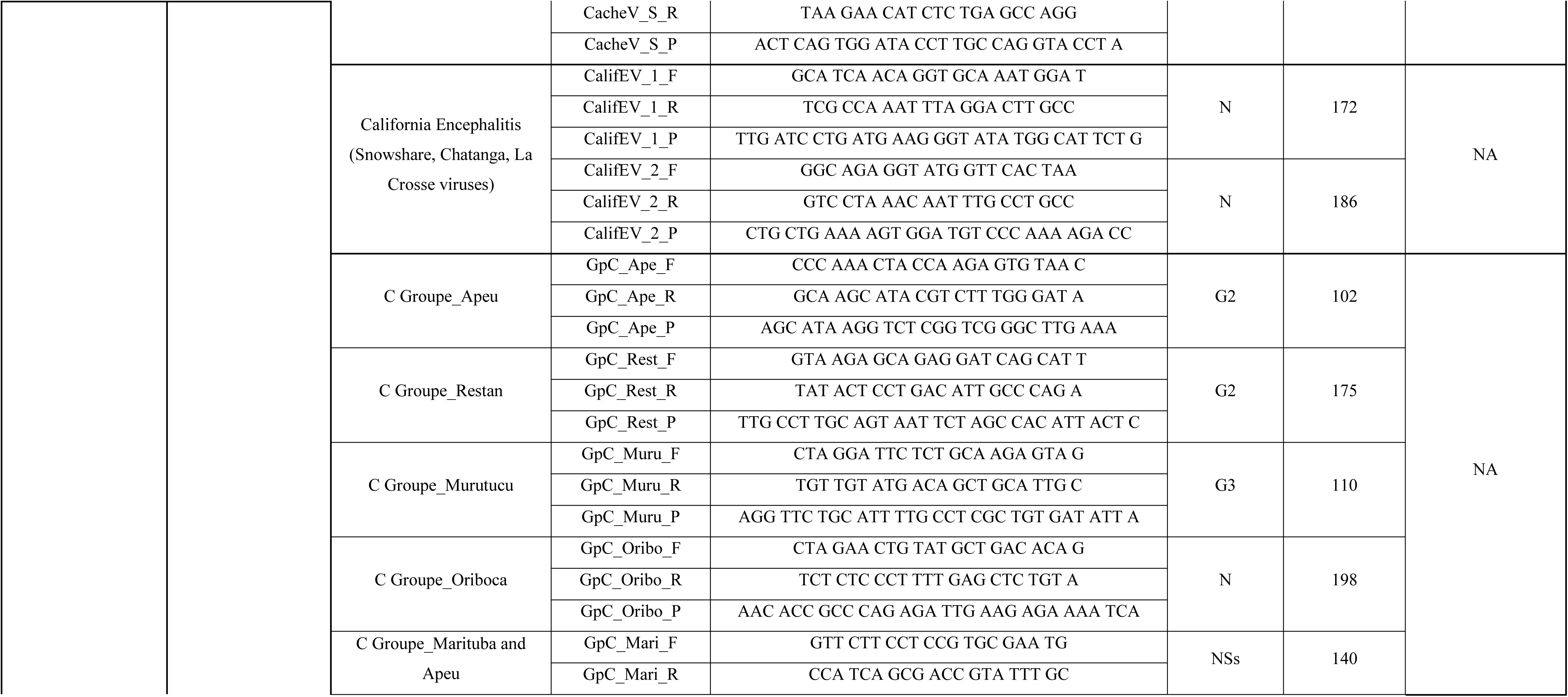
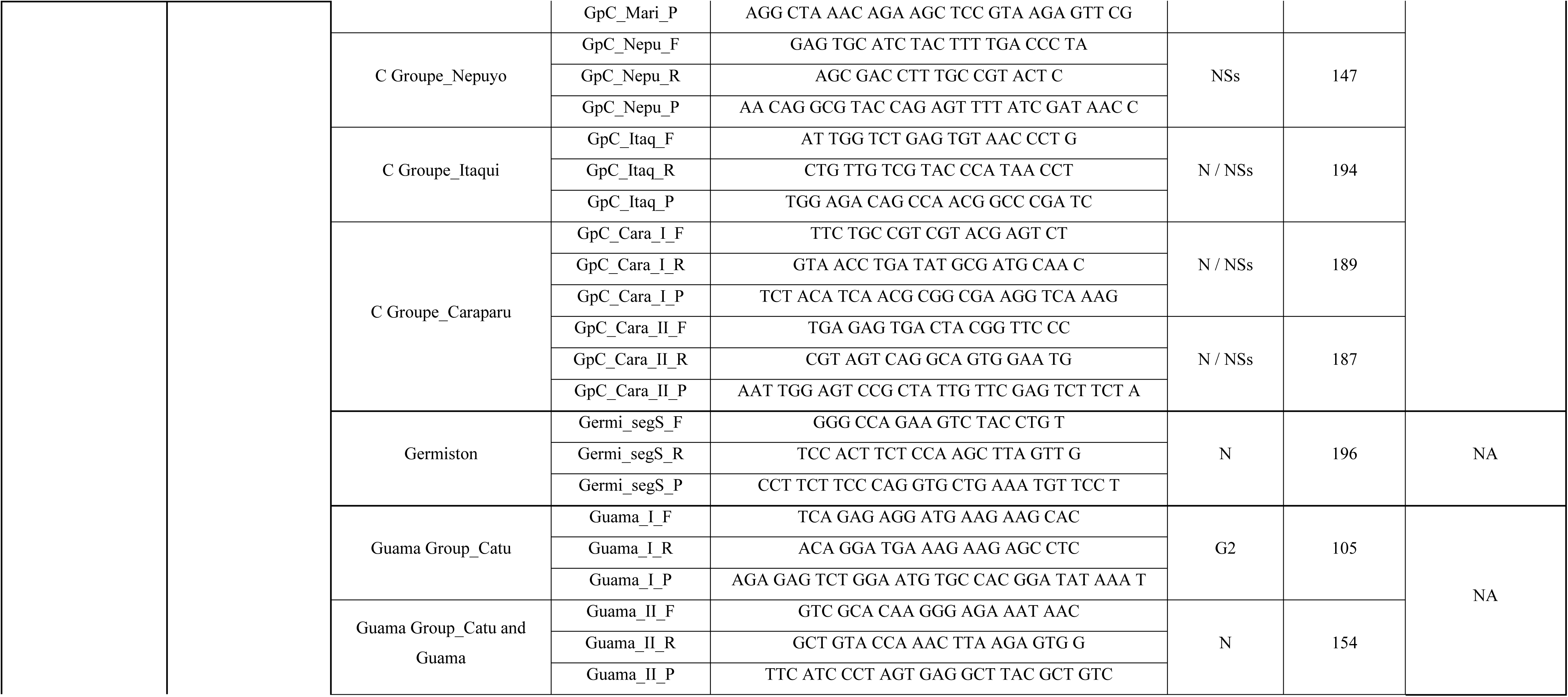
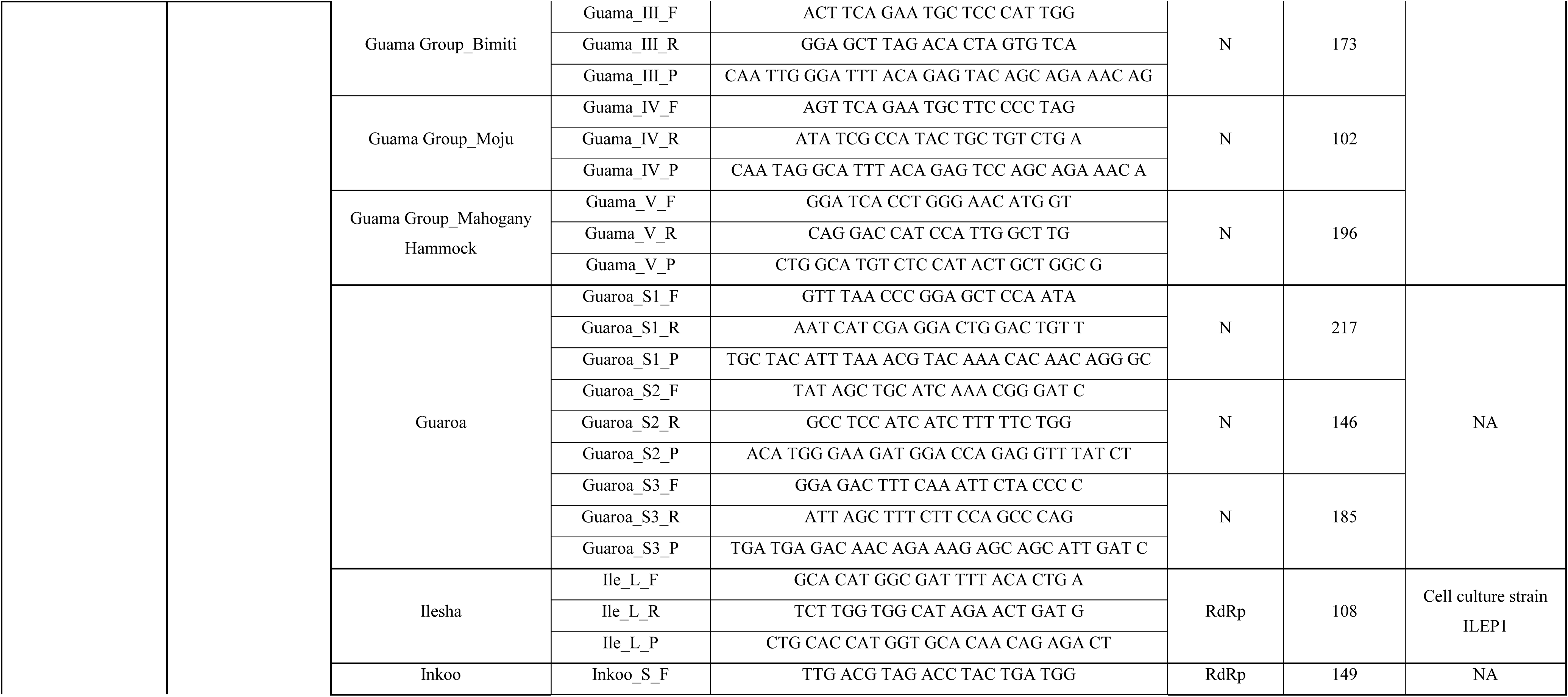
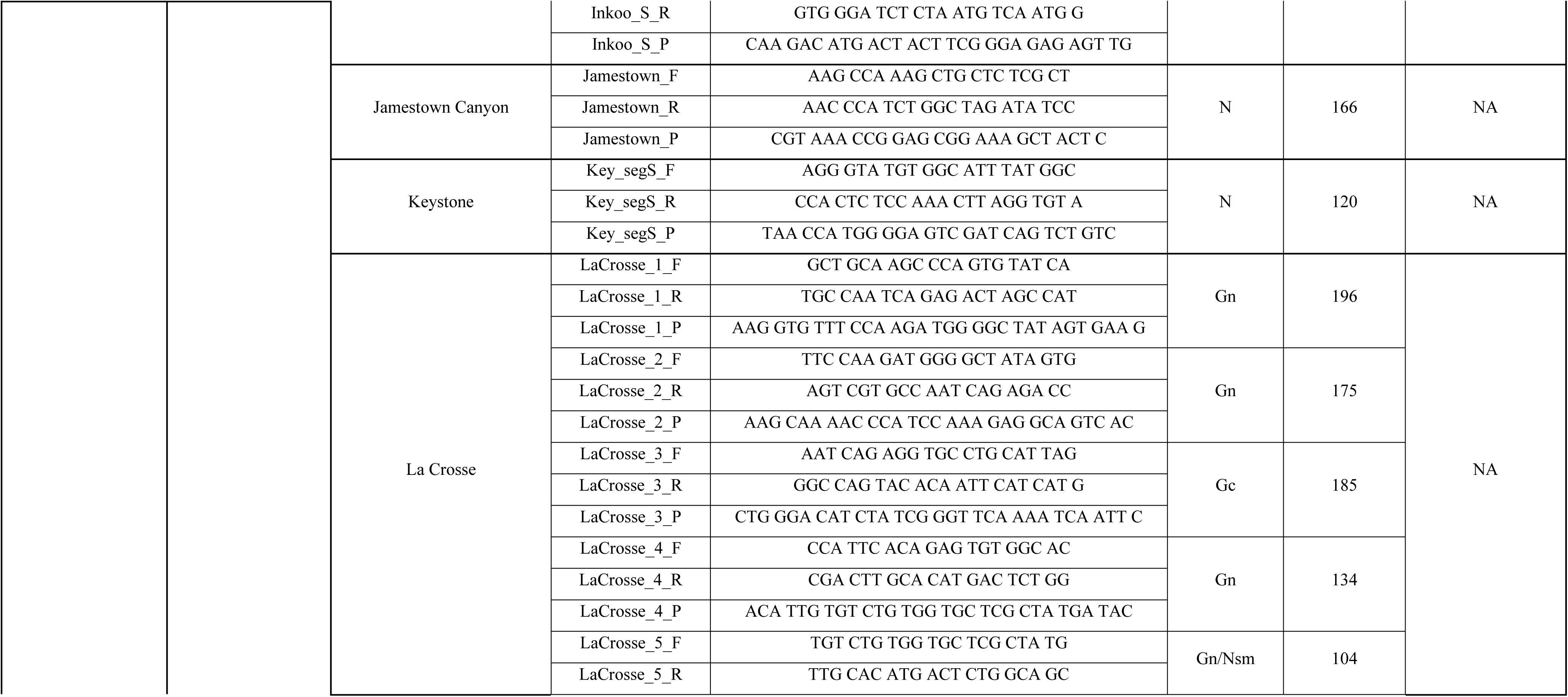
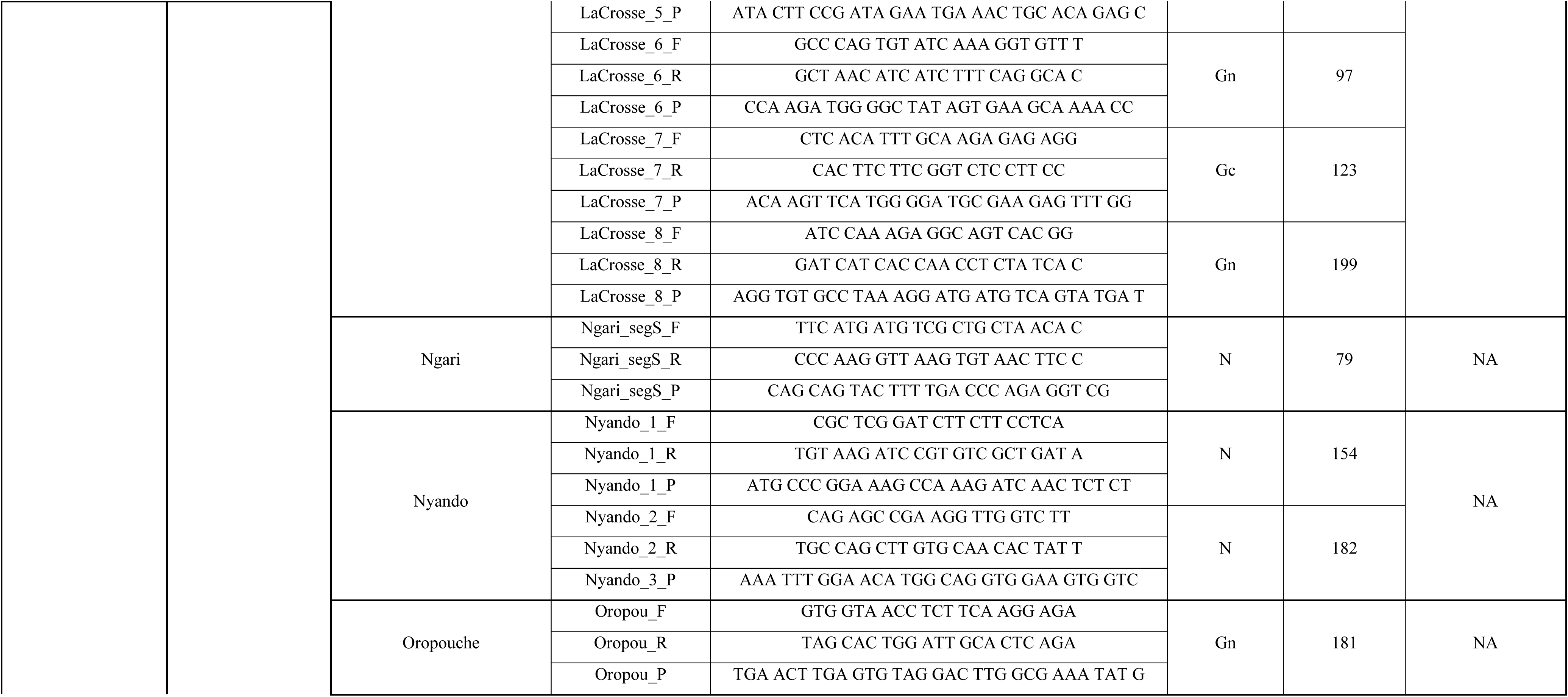
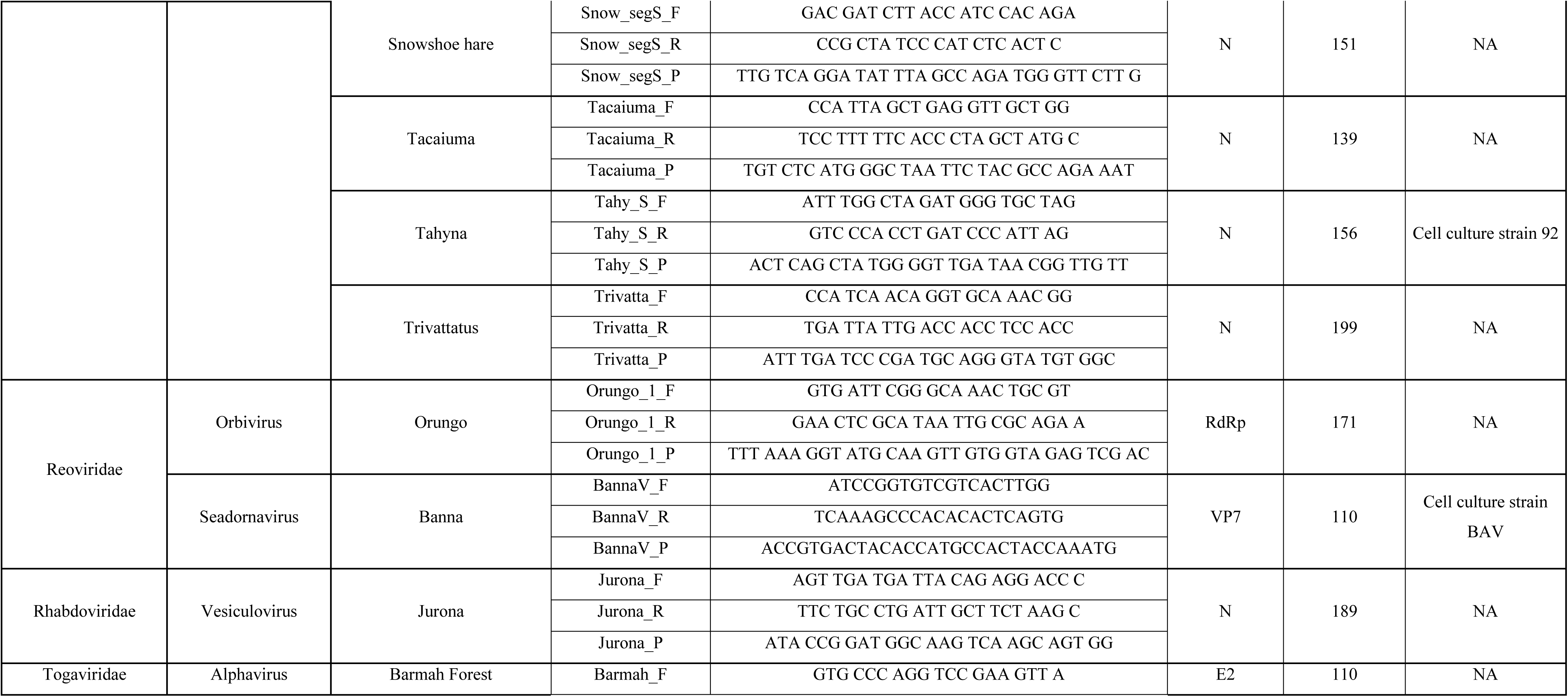
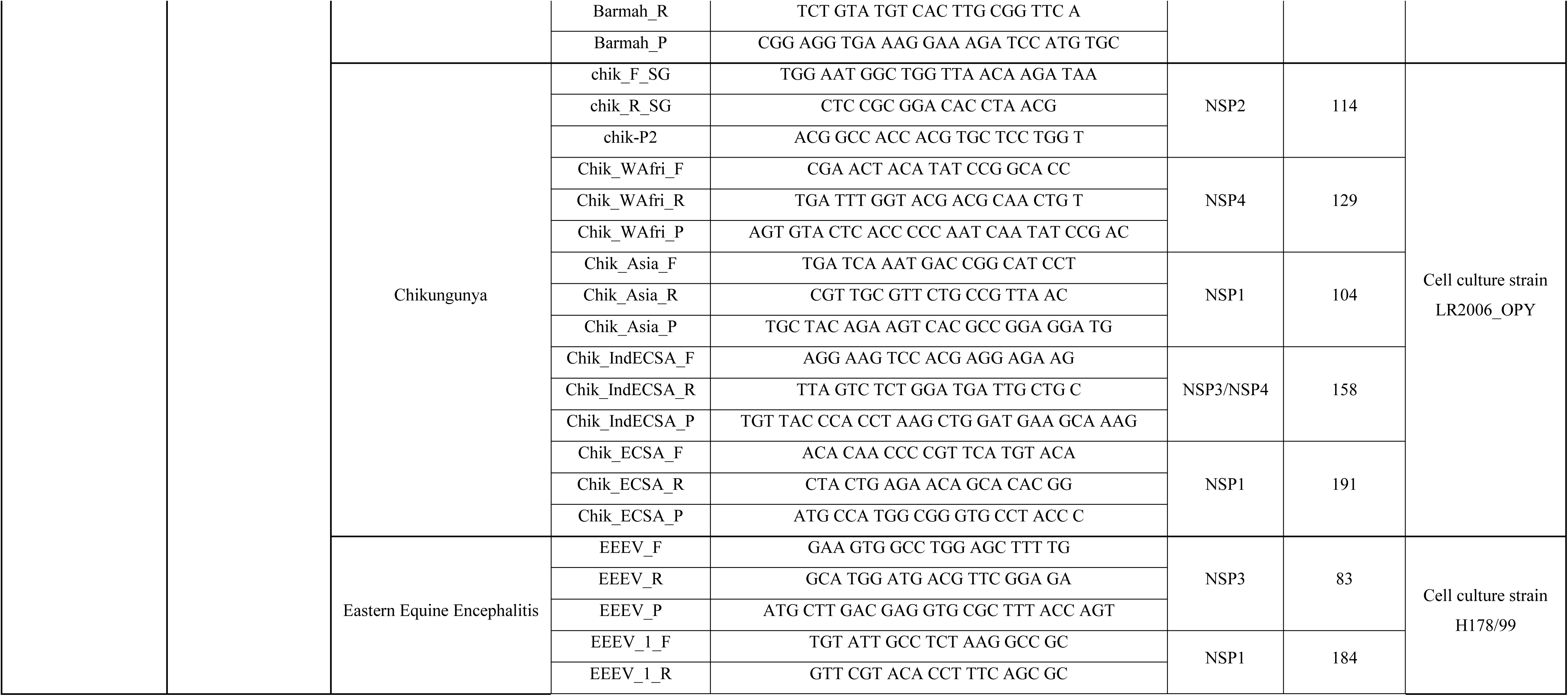
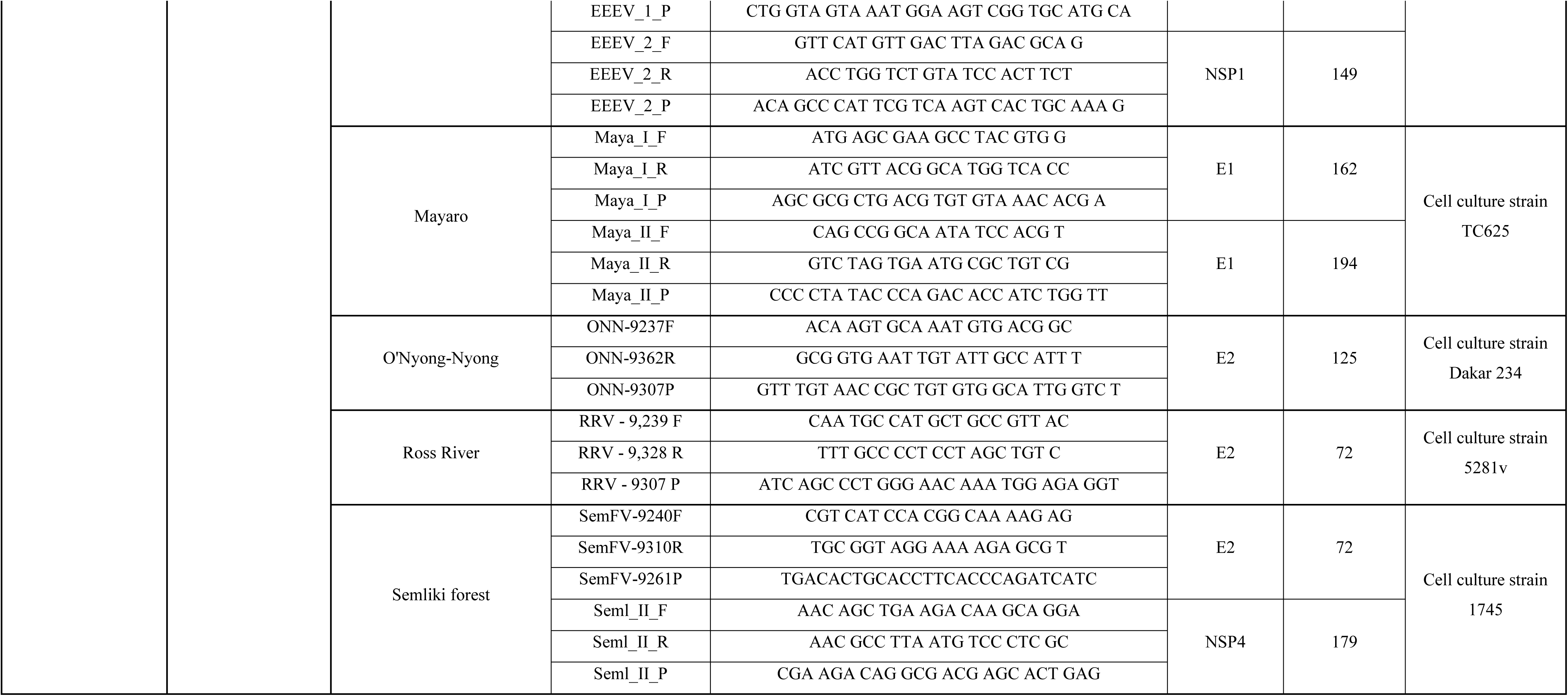
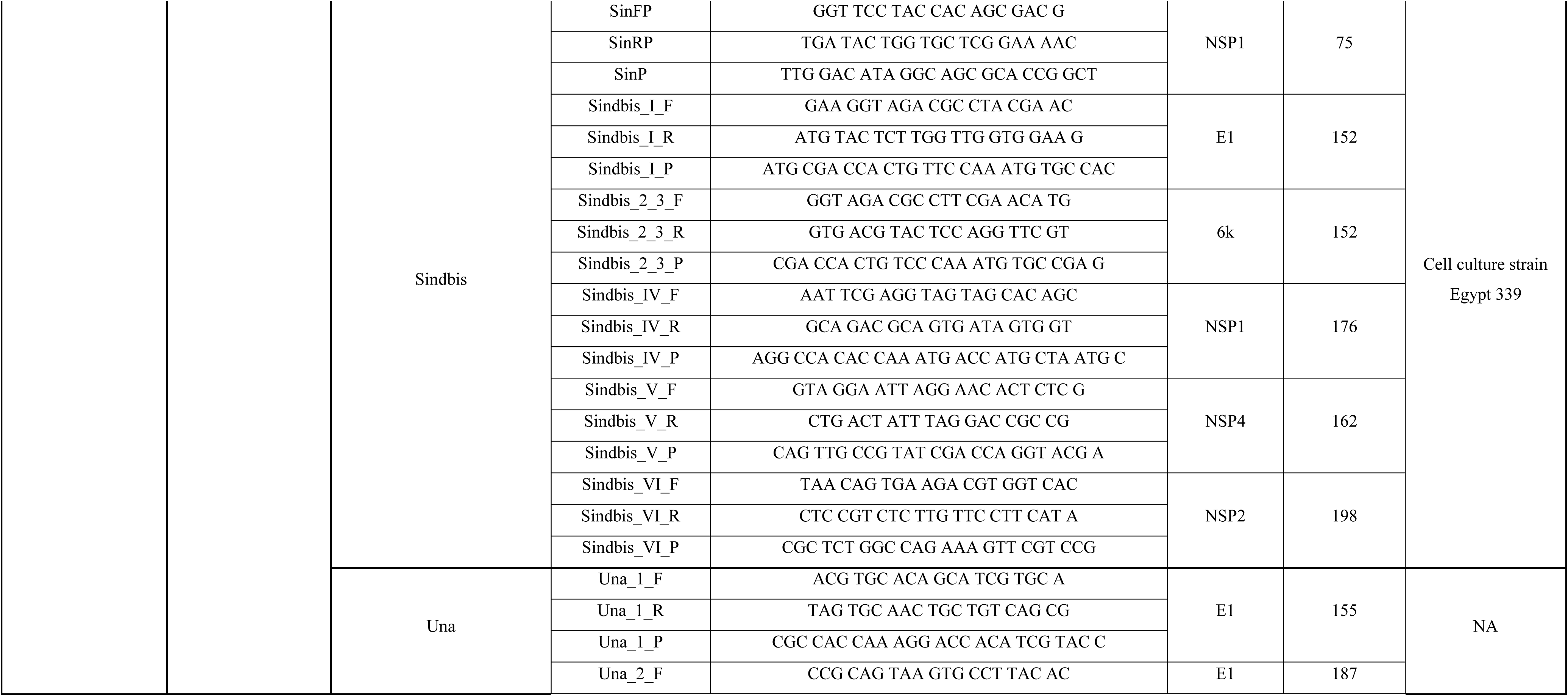
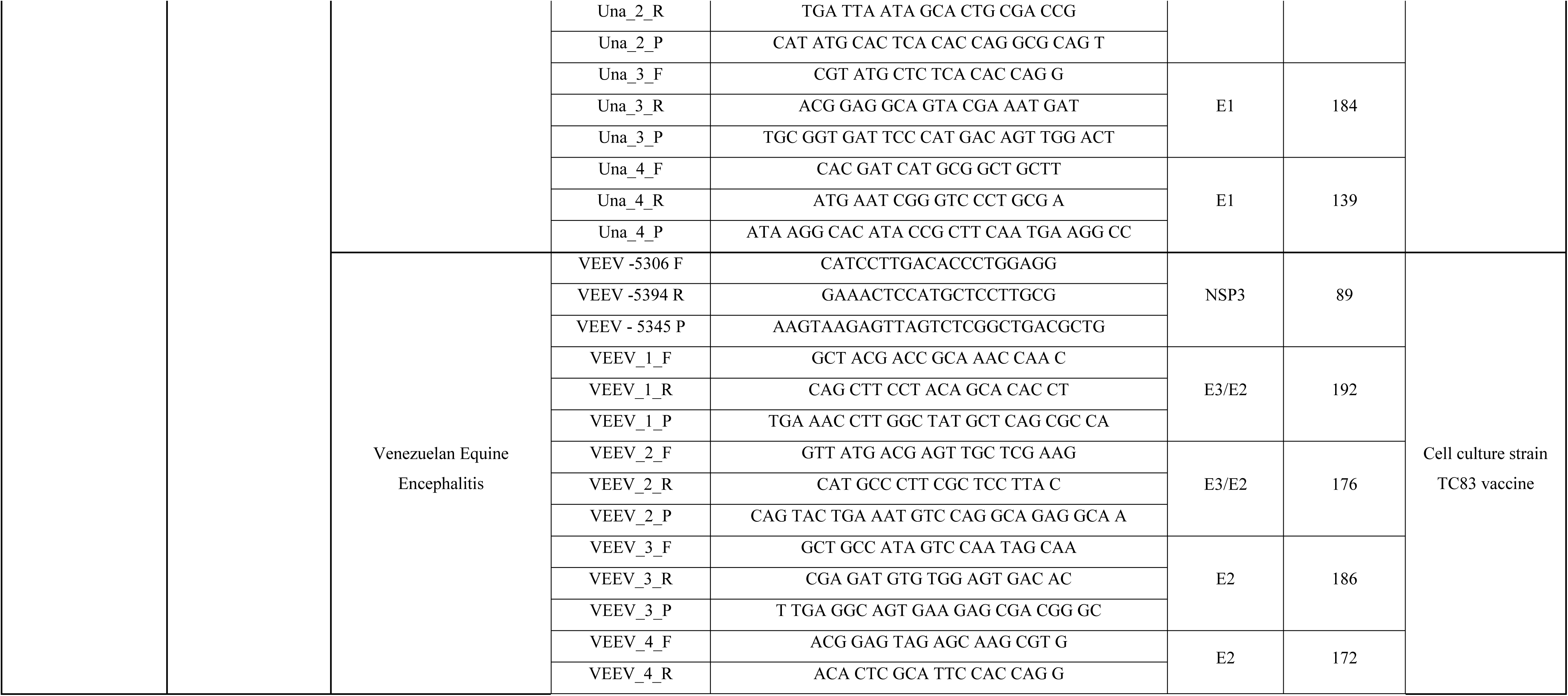
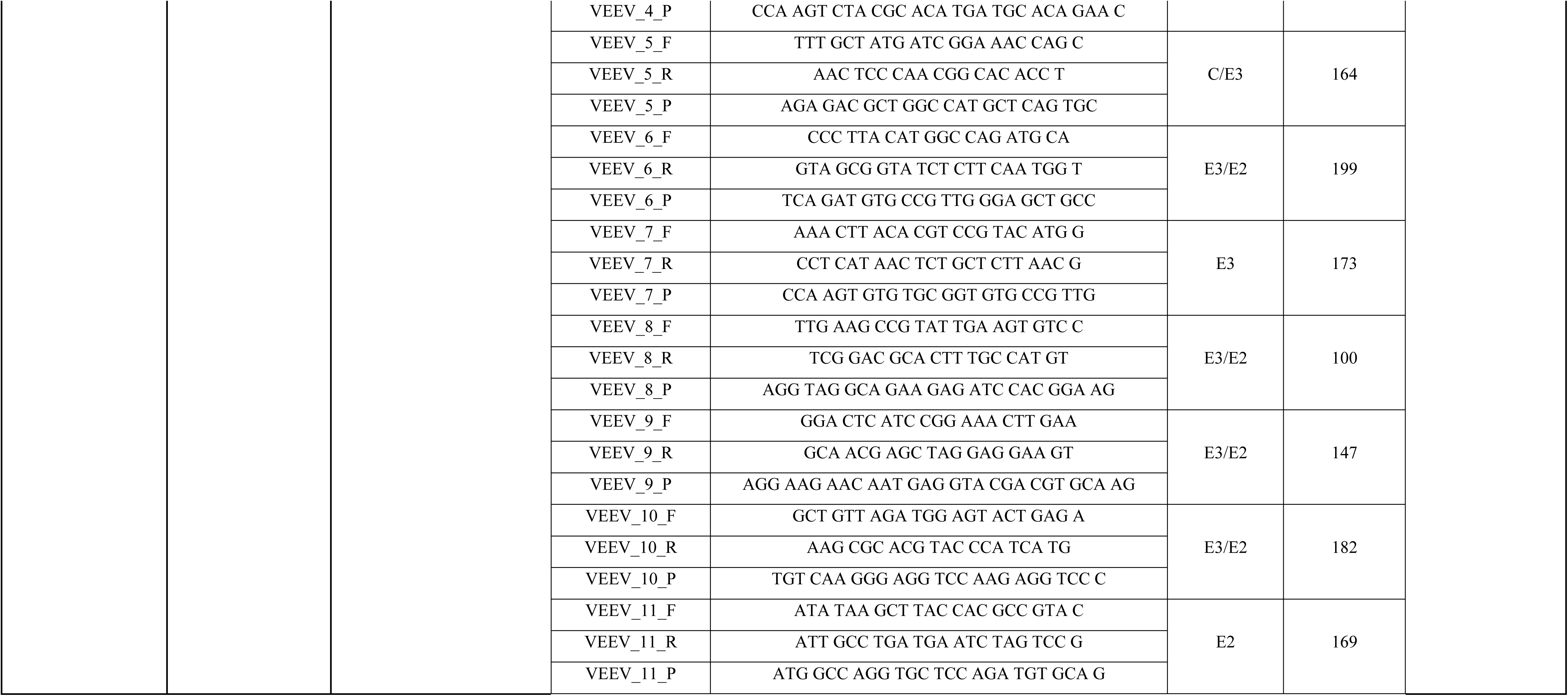
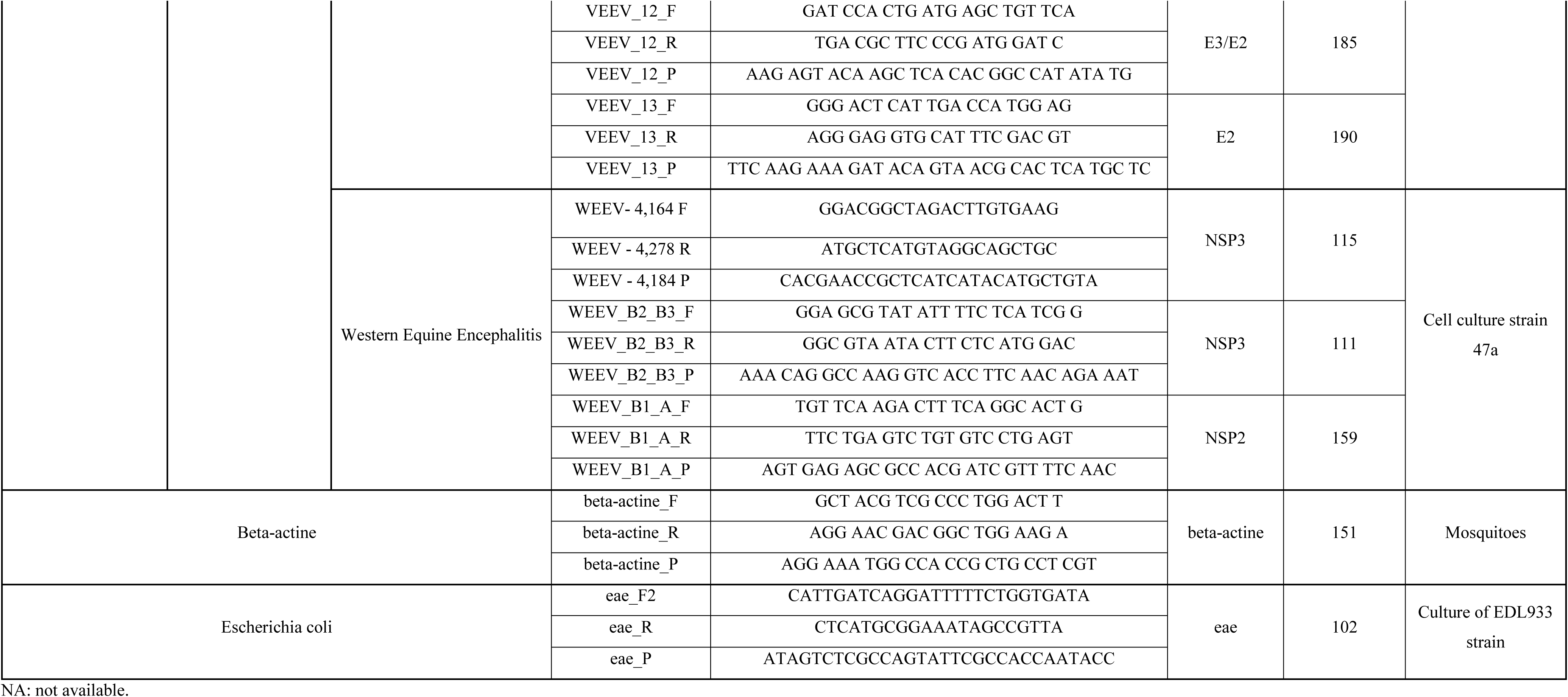
List of mosquito-borne viruses, targets, primers/probe sets, and positive controls.

**Table 8.**
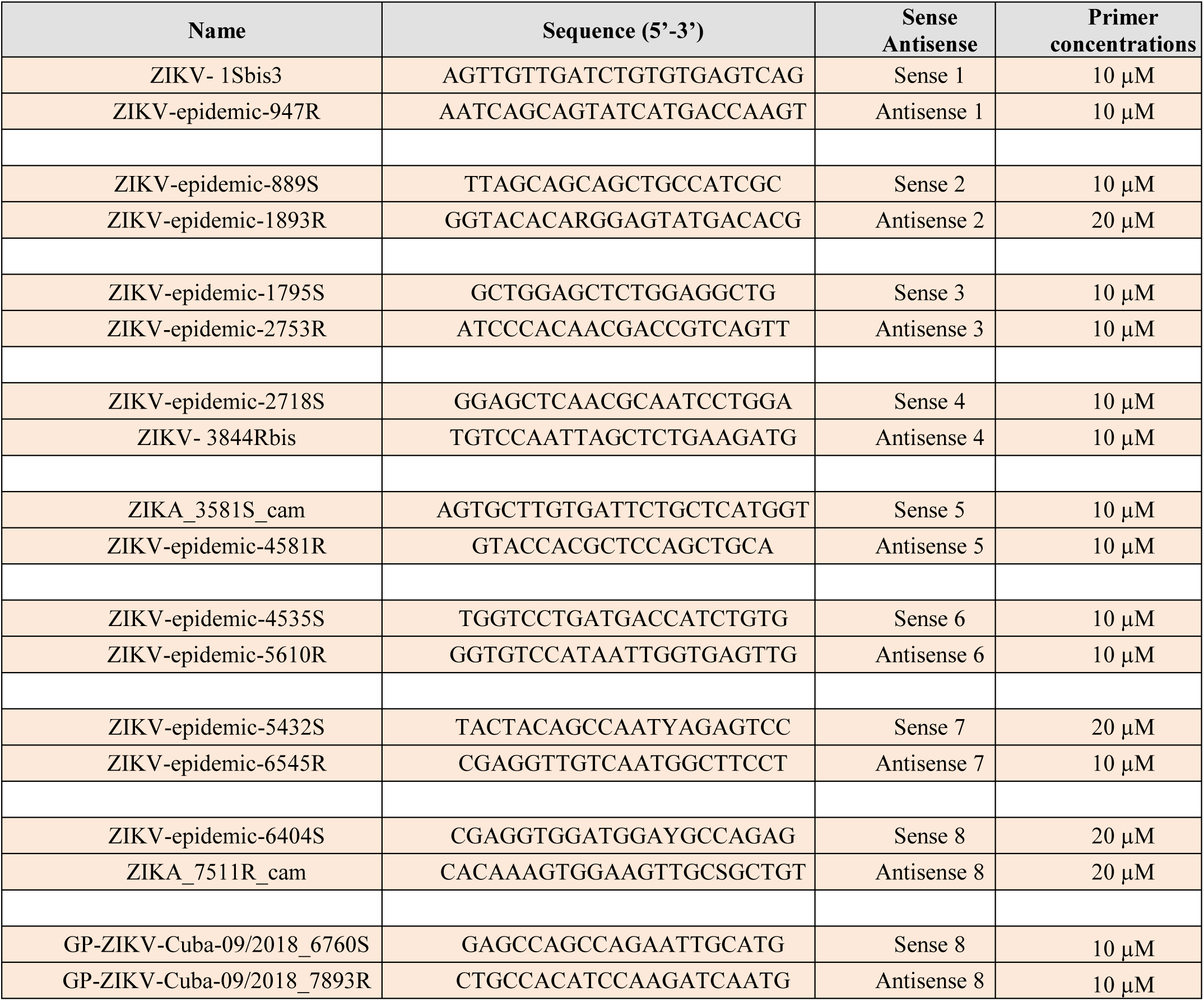
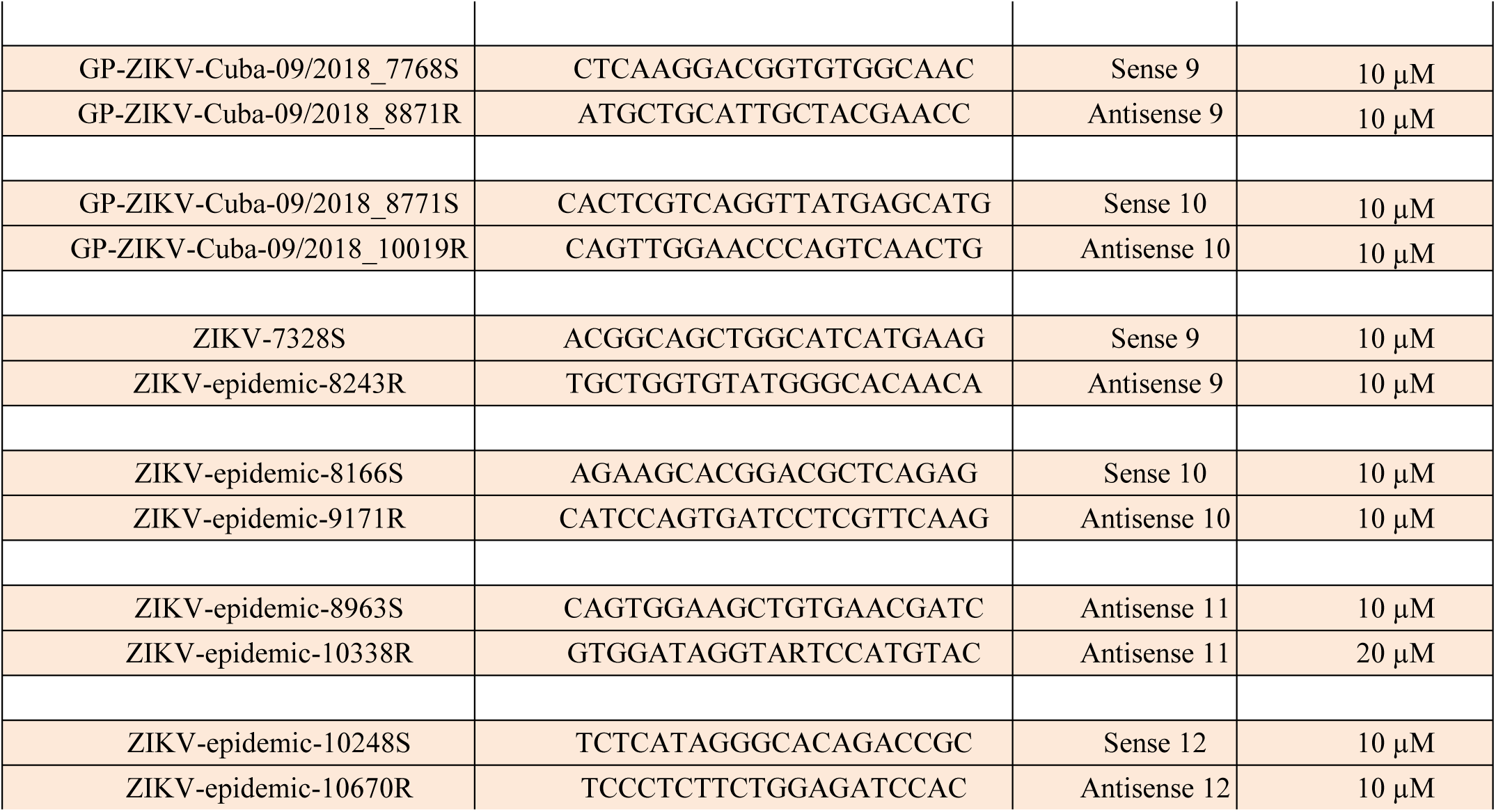
Primers used for full genome sequencing of Zika virus.

## 4. Discussion

In this study, we developed and validated a new high-throughput virus-detection assay based on microfluidic PCRs able to detect unambiguously 64 MBVs in mosquitoes. Only four primer sets demonstrated cross-reactivity with viruses from the same genus or serotype. Subsequently, we used this newly developed assay to perform a large epidemiological survey screening in six countries/territories during the last Zika pandemic. This new method has allowed the detection of (i) three human infecting arboviruses ZIKV, YFV and CHIKV in mosquitoes and (ii) other unexpected viruses such as TVTV.

The efficiency of our tool was the first requirement; we used artificially infected mosquitoes to detect different viruses (DENV1-4, CHIKV, WNV, ZIKV) offered in single and dual infections.

Our assays can target unambiguously DENV with however cross-reactions between serotypes. Caused by one of the four serotypes (DENV1-4), dengue is the most important arboviral disease worldwide (14). It is widely accepted that a subsequent infection with a second serotype can produce more severe symptoms (15). This situation becomes challenging when multiple serotypes co-circulate (16). Mosquitoes co-infected with different DENV serotypes are occasionally detected (17). *Aedes aegypti* and *Ae. albopictus* are urban vectors of DENV responsible for most epidemic outbreaks in Asia, Latin America, the Caribbean and Pacific islands (14). Co-infected *Aedes* mosquitoes are capable of transmitting multiple arboviruses during one bite (9). Dual DENV detections in mosquitoes may be a sign of co-circulation of DENV and then, may help in predicting co-infections in humans. Diagnosis of dengue infections cannot be based on clinical symptoms as dengue disease shares common symptoms with other arboviral diseases (18). To discriminate dengue serotypes, viral isolation and viral RNA detection remain the gold standard methods but should be performed during patient viremia (within five days after the onset of fever). Less constraining and costly, mass viral screening of mosquitoes in surveillance and epidemic contexts can be an advantageous substitute.

In the same way, WNV assays cross-reacted with the phylogenetically-related USUV. WNV is a flavivirus responsible of neuro-invasive disease in Europe and North America (19). Diagnosis of WNV infection remains challenging and human cases are usually underestimated. On the other hand, USUV has spread over Europe during the last twenty years causing bird mortalities and some rare human cases (20). Human infections are rare and often asymptomatic and neurological disorders can be described (20). While WNV circulates in Europe since 1960s, USUV shares the same geographical distribution and also the same vectors, *Culex pipiens*. Our tool did not succeed in distinguishing the two viruses and therefore, it needs more improvements.

By screening 17,958 mosquitoes collected in six countries/territories for 64 different MBVs, we succeeded in detecting ZIKV, YFV, CHIKV and TVTV in mosquitoes.

The Zika outbreak was unexpected; the first human cases outside endemic regions in Africa were reported in Yap island in 2007 where the outbreak was poorly publicized despite the two third of the population affected (21). Few years later, ZIKV hit French Polynesia (22) where the first notification of severe symptoms associated to ZIKV infections were done, Guillain-Barre syndrome (23) and microcephaly in new-born (24). After, ZIKV reached the American continent in 2015 (25), phylogenetic analysis indicated that the circulating ZIKV belonged to the Asian clade (26, 27). Our tool was able to detect ZIKV in pools of abdomen of *Ae. aegypti* and *Cx. pipiens* from Guadeloupe and French Guiana. However when analyzing disseminated viral particles in head and thorax, only *Ae. aegypti* was found infected corroborating the main role of this species in ZIKV transmission and excluding *Cx. quinquefasciatus* and *Cx. pipiens* as a vector (28).

Our mass screening tool has detected YFV in five mosquito species: *Aedes scapularis, Aedes taeniorhynchus, Haemagogus janthinomys, Haemagogus leucocelaenus,* and *Sabethes chloropterus.* The species *Hg. janthinomys* and *Hg. leucocelaenus* are considered as the main vectors of YFV in Brazil (29, 30) while *Aedes scapularis, Aedes taeniorhynchus* and *Sabethes chloropterus* only play a secondary role (31). Other viruses preliminary detected in Brazilian mosquitoes were TVTV and CHIKV. While CHIKV continues to cause sporadic cases in Brazil after the massive outbreak in 2015, TVTV was first isolated from *Aedes trivittatus* in USA in 1948, and has never been detected outside North America where it is mainly distributed (32). Consequences of TVTV infections on humans remain unknown (33). Nevertheless, the presence of CHIKV and TVTV in tested mosquitoes was not confirmed.

This study demonstrates the feasibility of high-throughput screening methods to detect diverse MBVs in field-collected mosquitoes. Performing 9,216 real-time PCRs in one run took four hours, and the cost was around $10 per reaction from sample homogenization to virus detection by real-time PCR (10, 11). Another main advantage of our tool is the adaptability of the system by adding new sets of primers and probes targeting newly emergent viruses in contrast to arrays with fixed panels of probes. Indeed, because the number of YFV cases was unusually high since January 2016 (34), we added specific detections of YFV strains circulating in South America to screen field-collected mosquitoes from Brazil, French Guiana, Suriname and Guadeloupe. In conclusion, our method designed to specifically identify MBVs in mosquitoes can be used to screen other types of samples such as human and/or animal blood or organs (35). We demonstrated the usefulness of this new screening method which represents a powerful, cost-effective and rapid system to track MBVs all around the world and could be easily customized to any viral emergence.

## Acknowledgements

We thank Christelle Delannay and the staff of the Regional Health Agency of Guadeloupe for their support during the sampling campaigns. This work was supported by the European Union’s Horizon 2020 Research and Innovation Programme under ZIKAlliance Grant Agreement no. 734548. The project also received funding from the 2014 PTR Anses-Institut Pasteur project (N° 511) for CHIPARBO, and the European Union’s Horizon 2020 Research and Innovation Programme under EVAg Grant Agreement no. 653316, and from an “Investissement d’Avenir” ‘grant from the Agence Nationale de la Recherche (CEBA ANR-10-LABX-2501 grant) supporting the project TIGERAMAZON.. The French General Directorate of Health funded the work performed in Guadeloupe and French Guiana. The Programme Opérationnel FEDER-Guadeloupe-Conseil Régional 2014–2020 (grant 2015-FED-192) supported the researchers from Guadeloupe. The authors declare they have no conflict of interest.

## 5. Author contributions

All the authors are involved as partners in the H2020 ZIKAlliance project. SM and ABF designed the experiments. SM and ABF wrote the paper. All the authors performed the experiments, and/or collected mosquitoes, and/or extracted RNAs, and/or performed confirmation tests like conventional and real-time PCR and virus isolation attempts. All the authors reviewed the manuscript.

## References

1. Bonaldo MC, Gomez MM, Dos Santos AA, Abreu FVS, Ferreira-de-Brito A, Miranda RM, et al. Genome analysis of yellow fever virus of the ongoing outbreak in Brazil reveals polymorphisms. Mem Inst Oswaldo Cruz. 2017;112(6):447–51.

2. Kramer LD, Ebel GD. Dynamics of flavivirus infection in mosquitoes. Advances in virus research. 2003;60:187–232.

3. Jupille H, Seixas G, Mousson L, Sousa CA, Failloux AB. Zika Virus, a New Threat for Europe? PLoS Negl Trop Dis. 2016;10(8):e0004901.

4. Weaver SC, Barrett AD. Transmission cycles, host range, evolution and emergence of arboviral disease. Nat Rev Microbiol. 2004;2(10):789–801.

5. Weaver SC, Charlier C, Vasilakis N, Lecuit M. Zika, Chikungunya, and Other Emerging Vector-Borne Viral Diseases. Annu Rev Med. 2018;69:395–408.

6. Dupont-Rouzeyrol M, O’Connor O, Calvez E, Daures M, John M, Grangeon JP, et al. Co-infection with Zika and dengue viruses in 2 patients, New Caledonia, 2014. Emerg Infect Dis. 2015;21(2):381–2.

7. Villamil-Gomez WE, Gonzalez-Camargo O, Rodriguez-Ayubi J, Zapata-Serpa D, Rodriguez-Morales AJ. Dengue, chikungunya and Zika co-infection in a patient from Colombia. Journal of infection and public health. 2016;9(5):684–6.

8. Caron M, Paupy C, Grard G, Becquart P, Mombo I, Nso BB, et al. Recent introduction and rapid dissemination of Chikungunya virus and Dengue virus serotype 2 associated with human and mosquito coinfections in Gabon, central Africa. Clin Infect Dis. 2012;55(6):e45–53.

9. Vogels CBF, Ruckert C, Cavany SM, Perkins TA, Ebel GD, Grubaugh ND. Arbovirus coinfection and co-transmission: A neglected public health concern? PLoS Biol. 2019;17(1):e3000130.

10. Michelet L, Delannoy S, Devillers E, Umhang G, Aspan A, Juremalm M, et al. High-throughput screening of tick-borne pathogens in Europe. Frontiers in cellular and infection microbiology. 2014;4:103.

11. Gondard M, Michelet L, Nisavanh A, Devillers E, Delannoy S, Fach P, et al. Prevalence of tick-borne viruses in Ixodes ricinus assessed by high-throughput real-time PCR. Pathog Dis. 2018;76(8).

12. Michelet L, Delannoy S, Devillers E, Umhang G, Aspan A, Juremalm M, et al. High-throughput screening of tick-borne pathogens in Europe. Front Cell Infect Microbiol. 2014;4:103.

13. Nielsen EM, Andersen MT. Detection and characterization of verocytotoxin-producing Escherichia coli by automated 5’ nuclease PCR assay. J Clin Microbiol. 2003;41(7):2884–93.

14. Bhatt S, Gething PW, Brady OJ, Messina JP, Farlow AW, Moyes CL, et al. The global distribution and burden of dengue. Nature. 2013;496(7446):504–7.

15. Katzelnick LC, Gresh L, Halloran ME, Mercado JC, Kuan G, Gordon A, et al. Antibody-dependent enhancement of severe dengue disease in humans. Science. 2017;358(6365):929–32.

16. Vaddadi K, Gandikota C, Jain PK, Prasad VSV, Venkataramana M. Co-circulation and co-infections of all dengue virus serotypes in Hyderabad, India 2014. Epidemiol Infect. 2017;145(12):2563–74.

17. Vazeille M, Gaborit P, Mousson L, Girod R, Failloux AB. Competitive advantage of a dengue 4 virus when co-infecting the mosquito *Aedes aegypti* with a dengue 1 virus. BMC Infect Dis. 2016;16:318.

18. Guzman MG, Harris E. Dengue. Lancet. 2015;385(9966):453–65.

19. Ciota AT, Kramer LD. Vector-virus interactions and transmission dynamics of West Nile virus. Viruses. 2013;5(12):3021–47.

20. Weissenbock H, Kolodziejek J, Fragner K, Kuhn R, Pfeffer M, Nowotny N. Usutu virus activity in Austria, 2001-2002. Microbes Infect. 2003;5(12):1132–6.

21. Duffy MR, Chen TH, Hancock WT, Powers AM, Kool JL, Lanciotti RS, et al. Zika virus outbreak on Yap Island, Federated States of Micronesia. N Engl J Med. 2009;360(24):2536–43.

22. Musso D, Nilles EJ, Cao-Lormeau VM. Rapid spread of emerging Zika virus in the Pacific area. Clinical Microbiology and Infection. 2014;20(10):O595–6.

23. Cao-Lormeau VM, Blake A, Mons S, Lastere S, Roche C, Vanhomwegen J, et al. Guillain-Barre Syndrome outbreak associated with Zika virus infection in French Polynesia: a case-control study. Lancet. 2016;387(10027):1531–9.

24. Cauchemez S, Besnard M, Bompard P, Dub T, Guillemette-Artur P, Eyrolle-Guignot D, et al. Association between Zika virus and microcephaly in French Polynesia, 2013-15: a retrospective study. Lancet. 2016;387(10033):2125–32.

25. Campos GS, Bandeira AC, Sardi SI. Zika Virus Outbreak, Bahia, Brazil. Emerg Infect Dis. 2015;21(10):1885–6.

26. Enfissi A, Codrington J, Roosblad J, Kazanji M, Rousset D. Zika virus genome from the Americas. Lancet. 2016;387(10015):227–8.

27. Zanluca C, Melo VC, Mosimann AL, Santos GI, Santos CN, Luz K. First report of autochthonous transmission of Zika virus in Brazil. Mem Inst Oswaldo Cruz. 2015;110(4):569–72.

28. Roundy CM, Azar SR, Brault AC, Ebel GD, Failloux AB, Fernandez-Salas I, et al. Lack of evidence for Zika virus transmission by Culex mosquitoes. Emerg Microbes Infect. 2017;6(10):e90.

29. Shannon RC, Whitman L, Franca M. Yellow Fever Virus in Jungle Mosquitoes. Science. 1938;88(2274):110–1.

30. Mascheretti M, Tengan CH, Sato HK, Suzuki A, de Souza RP, Maeda M, et al. [Yellow fever: reemerging in the state of Sao Paulo, Brazil, 2009]. Rev Saude Publica. 2013;47(5):881–9.

31. Abreu FVS, Ribeiro IP, Ferreira-de-Brito A, Santos A, Miranda RM, Bonelly IS, et al. Haemagogus leucocelaenus and Haemagogus janthinomys are the primary vectors in the major yellow fever outbreak in Brazil, 2016-2018. Emerg Microbes Infect. 2019;8(1):218–31.

32. Eklund C. Trivittatus virus. In: Karabatsos N, editor. International Catalogue of Arboviruses Including Certain Other Virus of Vertebrates. San Antonio, Texas: American Society of Tropical Medicine and Hygiene 1985. p. 1035–6.

33. Groseth A, Vine V, Weisend C, Ebihara H. Complete genome sequence of trivittatus virus. Arch Virol. 2015;160(10):2637–9.

34. Possas C, Martins RM, Oliveira RL, Homma A. Urgent call for action: avoiding spread and re-urbanisation of yellow fever in Brazil. Mem Inst Oswaldo Cruz. 2018;113(1):1–2.

35. Malmsten J, Dalin A, Moutailler S, Devillers E, Gondard M, Felton A, et al. Vector-Borne Zoonotic Pathogens in Eurasian Moose (Alces alces alces). Vector Borne Zoonotic Diseases. 2019;19(3):207–11.

